# The expanded *Bostrychia moritziana* genome unveils evolution in the most diverse and complex order of red algae

**DOI:** 10.1101/2025.02.11.637652

**Authors:** Romy Petroll, John A. West, Michael Ogden, Owen McGinley, Rory J. Craig, Susana M. Coelho, Michael Borg

## Abstract

Red seaweeds form part of an ancient lineage of eukaryotes that were one of the first to evolve multicellularity. Although they share a common evolutionary origin with modern-day plants and display complex multicellular development, we still lack comprehensive genome data from the most highly-evolved groups of red algae. Here, we present a chromosome-level genome assembly of *Bostrychia moritziana*, a species that is classified into the Rhodomelaceae family of the Ceramiales order, both of which constitute the largest and most diverse family and order of red algae, respectively. Contrary to the commonly held view that red algae generally have small genomes, we report significant genome size expansion in *Bostrychia* and other Ceramiales species, which we posit as one of at least three independent genome expansion events that occurred during red algal evolution. Our analyses suggest that these expansions do not involve polyploidy or ancient whole genome duplications, but in the case of *Bostrychia* appear to be largely driven by the dramatic proliferation of a single lineage of giant *Plavaka* DNA transposons. Consistent with increased genome size, we identify a substantial increase in gene content in *Bostrychia* that was shaped both by *de novo* gene emergence and by the amplification of gene families in common with other Ceramiales seaweeds, providing key insight into the genetic adaptations underpinning the evolutionary success of this species-rich order. Finally, our sex-specific assemblies enabled us to resolve the UV sex chromosomes in *Bostrychia*, which feature expanded gene-rich sex-linked regions. Notably, these sex-linked regions each harbour a distinct TALE-HD transcription factor orthologous to ancient regulators of haploid-diploid transitions in other multicellular lineages. Together, our findings offer a unique perspective of the genomic adaptations driving red algal diversity and demonstrate how this highly successful group of red seaweeds can provide insight into the evolutionary origins and universal principles of complex multicellular plant life.

## Introduction

The red algae are a highly diverse phylum of aquatic organisms in the Archaeplastida kingdom that share a common ancestor with modern-day land plants^1,2^. Fossil evidence suggests that the red algae were one of the first eukaryotic lineages to evolve multicellularity^3^, which is a remarkable feat considering that several genes normally essential in other eukaryotes were lost in their last common ancestor^4–6^. Although two independent transitions to multicellularity have been proposed during red algal evolution^7^, complex multicellularity is likely to have emerged only once in the Florideophyceae, one of seven Rhodophyta classes where the majority (∼95%) of species are classified (**Fig. 1A**). This dominant class of seaweeds display a diverse range of morphologies, ranging from simple filaments and blades to pseudoparenchymatous tissue and prominent reproductive structures in the most complex species^8,9^. This independent instance of complex multicellularity, coupled with its shared evolutionary origins with the green lineage, offers unique insight into the emergence of developmental complexity during Archaeplastida evolution.

**Figure 1.**
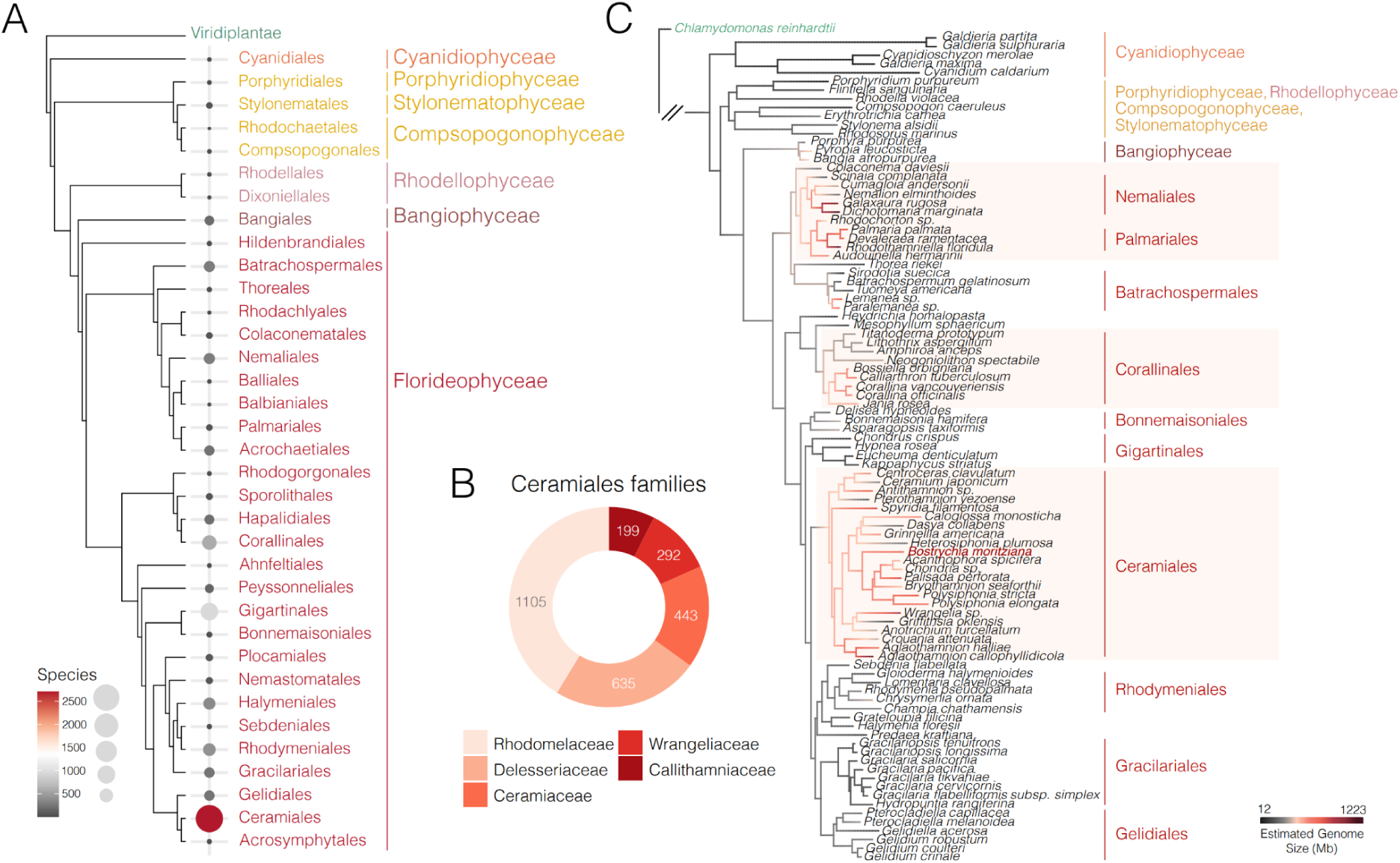
Species distribution and dynamics of genome size expansion across the red algae. (**A**) Phylogenetic tree showing all seven classes of red algae and their corresponding orders. Bubble sizes indicate the number of assigned species according to AlgaeBase^10^. (**B**) Pie chart illustrating the number of species assigned within the five families of the Ceramiales order. (**C**) Ancestral state reconstruction analysis using the genome size estimates of Kapraun & Freshwater (2012) as a continuous trait. Red shading highlights the origins of genome size expansion while the coloured lines on the tree indicate estimated genome sizes as detailed in the scale bar (in Mb).

Although the Florideophyceae is composed of several orders, the Ceramiales is by far the most diverse and species-rich, encompassing more than a third of all red algal taxa (**Fig. 1A**)^10^. The Ceramiales order has undergone significant diversification since its divergence from other red algal lineages around 390 million years ago, resulting in five distinct families with independent instances of emergent morphological complexity (**Fig. 1B**)^2,11^. An intriguing facet of the Ceramiales is their highly elaborate reproductive mechanisms, which have classically been used as distinguishing features in their taxonomic classification^11–13^. Like most of the Florideophyceae, the Ceramiales have a complex life cycle that alternates between three distinct multicellular generations: the haploid gametophytes, a diploid carposporophyte, and a diploid tetrasporophyte. While the free-living gametophytes and tetrasporophyte function similarly as in plants and other seaweeds (i.e., by producing gametes and meiotic spores, respectively), the carposporophyte is unique to the florideophycean red algae and serves to replicate the progeny into an abundant mass of spores, maximizing the success of each fertilization event^14^. This unique reproductive strategy, combined with the increased morphological complexity and broad diversification of the Ceramiales over 250 million years^2^, suggests the emergence of key innovations that are likely reflected in the evolution of their genomes. The Ceramiales are thus of particular interest for comparative genomic investigation, particularly given the lack of high-quality, sex-specific chromosome-level assemblies in this order.

## Results and discussion

### Red algal evolution involved at least three independent genome size expansion events

Theory has it that the common red algal ancestor experienced an evolutionary bottleneck that led to the purging of genomic content, leading to the loss of conserved traits such as flagellated stages, autophagy, and phytochrome-based light sensing^6,8,15^. Most red algal genomes studied thus far are generally small (∼100 Mb) with a low gene and intron count compared to other lineages^8^. However, a survey of nuclear DNA content in several red algal isolates has suggested substantial genome size variation across the Rhodophyta, with large genome sizes predicted in particular orders of the Florideophyceae^16^. Although these genome size estimates may not be exact, their relative proportions align with genomic sequencing data, including recent draft assemblies from several red algal orders^17,18^, with the smallest genome sizes occurring in the Cyanidiophyceae and the largest in the Florideophyceae (**Supplemental Fig. 1**; **Supplemental Table 1**).

To investigate the dynamics of genomic size expansion (GSE) during red algal evolution, we performed an ancestral state reconstruction (ASR) analysis using the genome size estimates of Kapraun & Freshwater (2012) as a continuous trait (**Fig. 1C**). For this, we first constructed a maximum likelihood tree based on at least three of seven conserved marker genes from 939 red algal and Viridiplantae species, using cryptophyte algae as an outgroup (**Supplemental Fig. 2**). Likelihood-based reconstruction of trait evolution using a pruned phylogeny resolved at least three independent origins of GSE within the Florideophyceae, specifically within the order Corallinales, within the subclass Nemaliophycidae, and at the node common to the Ceramiales (**Fig. 1C**). Flow cytometry analysis of several members of the Ceramiales confirmed large genome sizes that ranged between 1.22 and 3.64 Gb, which included four different species in the *Bostrychia* genus, *Polysiphonia senticulosa* and *Caloglossa monosticha* (**Supplemental Fig. 3**; **Supplemental Table 2**). GSE thus appears to be a common feature of the Ceramiales, which we hypothesised to have influenced the emergence of morphological complexity and evolutionary radiation of this highly successful order.

### Chromosome-level assembly of the *Bostrychia moritziana* genome

To investigate the molecular drivers of GSE in the Ceramiales, we focused our attention on the red alga *Bostrychia moritziana* (Sonder ex Kützing) (herein *Bostrychia*) (**Fig. 2A-D**)^19^. This filamentous epiphytic seaweed is common in mangrove habitats and is classified into the dominant Rhodomelaceae family of the Ceramiales order (**Fig. 1B**). *Bostrychia* has been subject to an important body of work characterising several aspects of its biology^20^, including its physiology^21^, life history^22,23^, reproduction^24–27^, ecology^21,28,29^ and genetic diversity^20,28–31^. We combined Oxford Nanopore and PacBio HiFi sequencing to generate draft genome assemblies of haploid male and female *Bostrychia* gametophytes, both siblings of a previously described South African isolate (**Fig. 2A-D**; **Supplemental Table 3**)^31^. The initial draft assemblies were both approximately 1.2 Gb long with 78.9% and 78.4% BUSCO completeness scores for the male and female, respectively **(Supplemental Table 3**).

**Figure 2.**
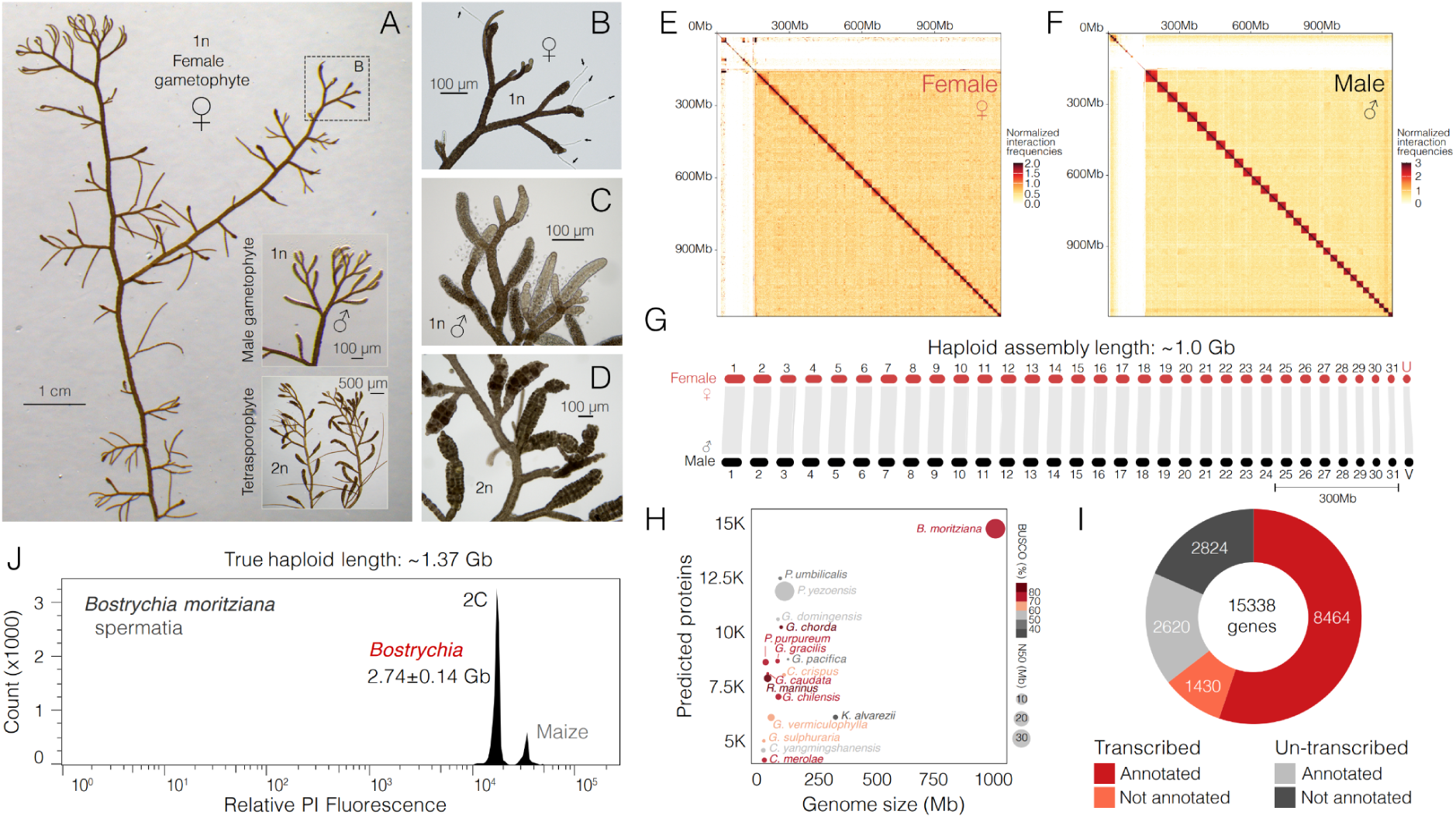
Sex-specific chromosome-level assembly of the *Bostrychia moritziana* genome. (**A**) A branch of a female *Bostrychia* gametophyte thallus is shown, with tips of the male gametophyte and tetrasporophyte (shown inset). (**B**) Close-up view of the female gametophyte tip in panel A. Mature carpogonial branches with protruding trichogynes as indicated by the black arrows. (**C**) Tips of a male gametophyte with mature spermatangia actively releasing spermatia. (**D**) Tetrasporangial stichidia of the tetrasporophyte loaded with maturing tetraspores. (**E**) Normalized Hi-C contact map showing interaction frequencies within the female assembly. (**F**) Normalized Hi-C contact map showing interaction frequencies within the male assembly. (**G**) Gene synteny plot comparing the male and female assemblies of the *Bostrychia* genome. Chromosome numbers are labeled while the sex chromosomes are indicated with U for the female and V for the male. (**H**) Bubble chart illustrating the number of predicted proteins plotted against genome size (in Mb) of several red algal species. Bubble size represents the N50 value (in Mb) while the colors indicate the BUSCO score of each genome assembly. (**I**) Pie chart summarising the number of transcribed and functionally annotated genes in *Bostrychia.* (**J**) Flow cytometry analysis of propidium iodide-stained spermatial nuclei from *Bostrychia* alongside nuclei isolated from leaf tissue of *Zea mays* (L. ‘CE-777’). The estimated DNA content of spermatial nuclei is indicated alongside the diploid (2C) peak, with the corresponding haploid genome size indicated above.

To resolve the assemblies at a chromosome-level, we integrated Hi-C data derived from the same male and female gametophyte strains and implemented an additional gap-closing strategy. This yielded 32 chromosomes in each assembly that were evident as distinct territories with strong intra-chromosomal interactions along the diagonal of Hi-C chromatin contact maps (**Fig. 2E&F**). Hi-C integration also helped purge contigs belonging to the substantial microbial community present within the cultures (140 Mb and 147 Mb in the male and female, respectively). A small percentage of unplaced contigs with inter-chromosomal interactions were also evident (21 Mb in the male and 27 Mb in the female), which likely represent difficult-to-resolve repeats (**Fig. 2E&F**). Importantly, we observed strong synteny across all 32 chromosomes between the male and female, verifying the robustness of our assembly (**Fig. 2G**). Telomeric sequences (TTTTTAGGGG) were identified on one end of 14 different chromosomes, while ribosomal RNA (rRNA) genes were present across the genome, with a large cluster prominent on chromosome 22 (**Supplemental Fig. 4**). With an assembly size of approximately 1.0 Gb, the *Bostrychia* genome is among the largest red algal genomes sequenced to date, validating predictions of increased genome size within the Ceramiales (**Fig. 2G&H**; **Table 1**). In addition to the nuclear genome, we also assembled and annotated the complete chloroplast and mitochondrial genomes (**Supplemental Fig. 5A&B**).

**Table 1.**
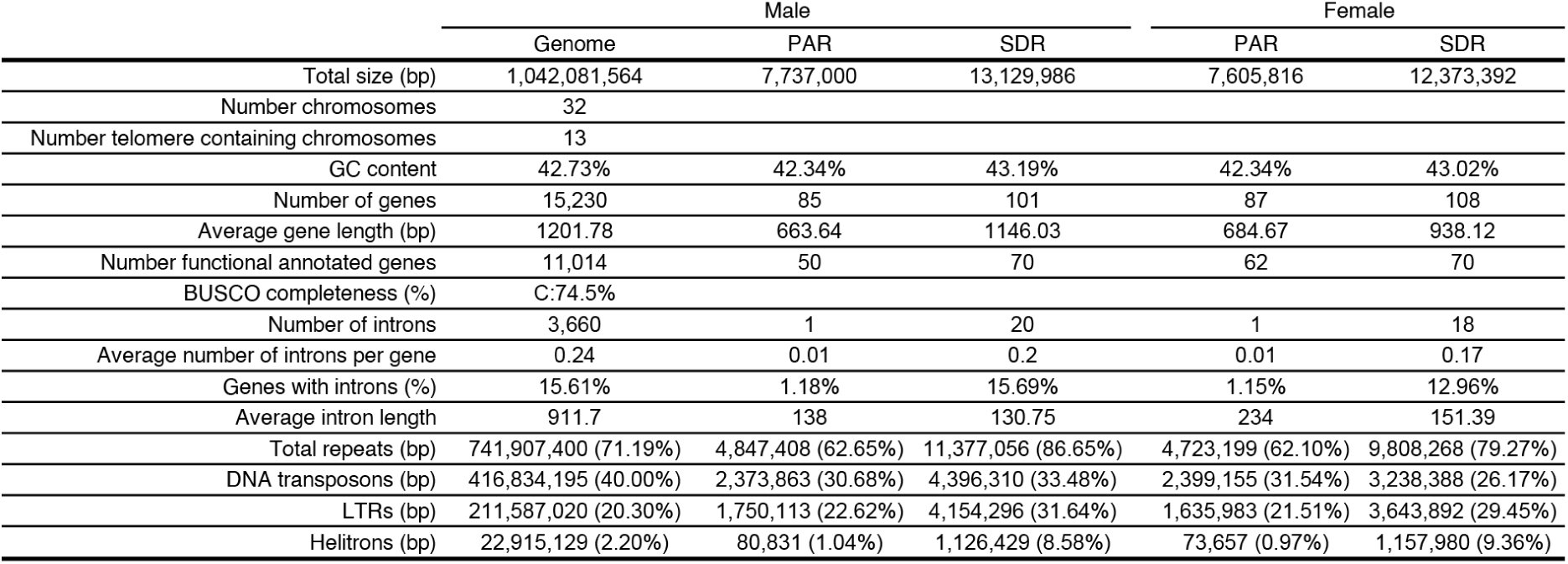
Summary statistics of the genome assembly of *Bostrychia moritziana*.

We predicted 15,338 gene models in *Bostrychia*, two-thirds of which (9,894; 64.5%) were supported by transcript evidence in at least one of the three major life stages we profiled (**Fig. 2I**; **Table 1; Supplemental Table 4&5**; **Supplemental Fig. 6A-C**). Most transcribed genes (8,464; 55.2%) had at least one domain present in the InterPro, eggNOG, or TAPscan databases, while 1,430 (9.3%) were specific to red algae and had no functional annotation (**Fig. 2I**; **Supplemental Table 5**). The remaining third were not expressed and were evenly distributed between annotated (2,620) and unannotated (2,824) genes (**Fig. 2I**). These untranscribed genes were predominantly young and specific to red algae or *Bostrychia,* mostly monoexonic, and significantly shorter compared to transcribed genes (**Supplemental Fig. 7A&B**). We speculate that these genes may be transcribed under specific developmental stages or environmental conditions and/or perhaps serve as a reservoir of “proto-genes” that might facilitate *de novo* gene birth^32^. As is typical of the more compact red algal genomes, the percentage of intron-containing genes was also low in *Bostrychia* (15.59%), although intron length was subtly and significantly increased compared to that of Gracilariales genomes and similar in lengths to that reported in *Chondrus* (Gigartinales) (**Supplemental Fig. 7C-F**). Thus, consistent with its dramatically increased size, the *Bostrychia* genome encodes an expanded gene repertoire when compared to other red algal orders (**Fig. 2H**; **Supplemental Fig. 7C-F**; **Supplemental Fig. 8**).

### Whole genome duplication is unlikely to have contributed to evolution of the Rhodomelaceae

The expanded genome and gene content of *Bostrychia* prompted us to investigate the molecular drivers underlying GSE. In land plants, polyploidy is highly prevalent and has significantly contributed to genome complexity^33,34^. Cytogenetic investigations have suggested that both aneuploidy and polyploidy might have accompanied speciation in the Ceramiales order^12,35–37^. We thus employed a series of genomic and phylogenomic approaches to clarify the potential impact of polyploidy in *Bostrychia* and the Ceramiales more generally.

We first performed flow cytometry analysis to verify the size of the *Bostrychia* genome assembly (**Supplemental Table 2**). In closely-related species of *Bostrychia* and *Polysiphonia* (Rhodomelaceae), spermatia and tetraspores are consistently released with a replicated haploid genome or 2C level of DNA^36,38,39^. Freshly-released spermatia and tetraspores of *Bostrychia* both contained 2.8 pg of DNA, indicating a genome size of around 2.7±0.1 Gb (**Fig. 2J**; **Supplemental Fig. 9A**). The haploid genome size of *Bostrychia* is thus likely closer to 1.35 Gb in length, probably due to unresolved repeats evident as orphan contigs with strong inter-chromosomal interactions in our Hi-C maps (**Fig. 2E&F**). Next, we measured DNA content in each life stage to assess ploidy changes across the *Bostrychia* life cycle. In the Rhodomelaceae, new cell growth emanates from a polyploid uninucleate apical cell^38,39^. In the male gametophyte of *Bostrychia radicans*, the apical cell has a 4C DNA content that is reduced to minimum of 2C in most cells of the mature thallus^38^. Consistently, both the male and female *Bostrychia* gametophyte contained the equivalent of 2C levels of DNA (**Supplemental Fig. 9B-C**), while the tetrasporophyte contained both 2C and 4C peaks that correspond to haploid tetraspores and diploid vegetative cells, respectively (**Supplemental Fig. 9D**). Our findings in *Bostrychia* thus corroborate earlier observations that most cells of the gametophyte and tetrasporophyte phases in the Rhodomelaceae are endo-diploid and endo-tetraploid, respectively. This has significant implications for red algal evolution and ecology, particularly with regard to how endopolyploidy may help buffer against deleterious mutations and maintain cryptic variation, thereby facilitating environmental adaptation and evolutionary change, which is also common in flowering plants^40^.

Next, we assessed patterns of sequence divergence to look for signs of a potential ancient whole genome duplication (WGD). The distribution of synonymous distances (*K*_s_) between syntenic paralogues in *Bostrychia* did not reveal a deviating retention rate indicative of a WGD event (**Supplemental Fig. 10A**). This was further reflected in the lack of inter-chromosomal synteny in each assembly as well as in the mostly constant gene family sizes *Bostrychia* shares with species from other red algal orders (**Supplemental Fig. 10B-D**). Fossil-calibrated phylogenies have estimated that, like the Ceramiales, the fern lineage of vascular plants emerged in the Paleozoic era *circa* 350-400 million years ago (mya)^2,41,42^. Given that ferns experienced at least two rounds of WGD in their evolutionary history, with the oldest occurring around 250 mya^43,44^, it is unlikely that evolutionary age precluded us from detecting traces of WGD in *Bostrychia*. We thus conclude that WGD is unlikely to have occurred since the emergence of Rhodomelaceae 250 mya and subsequently during evolution of the *Bostrychia* lineage. Nevertheless, we cannot rule out a deeply ancient and untraceable event having occurred at the dawn of the Ceramiales.

### Ancestral bursts in transposon proliferation underlie genome expansion in *Bostrychia*

Having ruled out polyploidy in *Bostrychia*, we turned our attention to the repeat complement in the genome. Repetitive elements overwhelmingly dominate the genomic landscape, accounting for a total of 744 Mb (71%), with more than a third (417 Mbp; ∼40%) classified as DD(E/D) DNA transposons (**Fig. 3A**; **Supplemental Fig. 8**). LTR elements are also abundant (213 Mbp (∼20%)), with LINEs, Cryptons and Helitrons present in a minor contribution (**Fig. 3A**). The distribution of TE divergence in *Bostrychia* suggests that the genome experienced at least two major bursts of TE proliferation (**Fig. 3B**). The first and oldest was primarily driven by LTR elements, while the second was overwhelmingly dominated by the proliferation of DNA transposons (**Fig. 3B**). Strikingly, the overwhelming majority of DNA transposons were represented by *Plavaka* elements, which belong to a superfamily of DNA transposons called *EnSpm* or *CACTA* first described in basidiomycete fungi (**Supplemental Fig. 11&12**)^45^. *Plavaka* elements alone account for just over a third of the genome (373 Mb; ∼36%), almost all of which is contributed by only three giant families (17.4 – 19.1 kb), highlighting their proliferation as a significant cause for GSE in *Bostrychia* (**Fig. 3A**).

**Figure 3.**
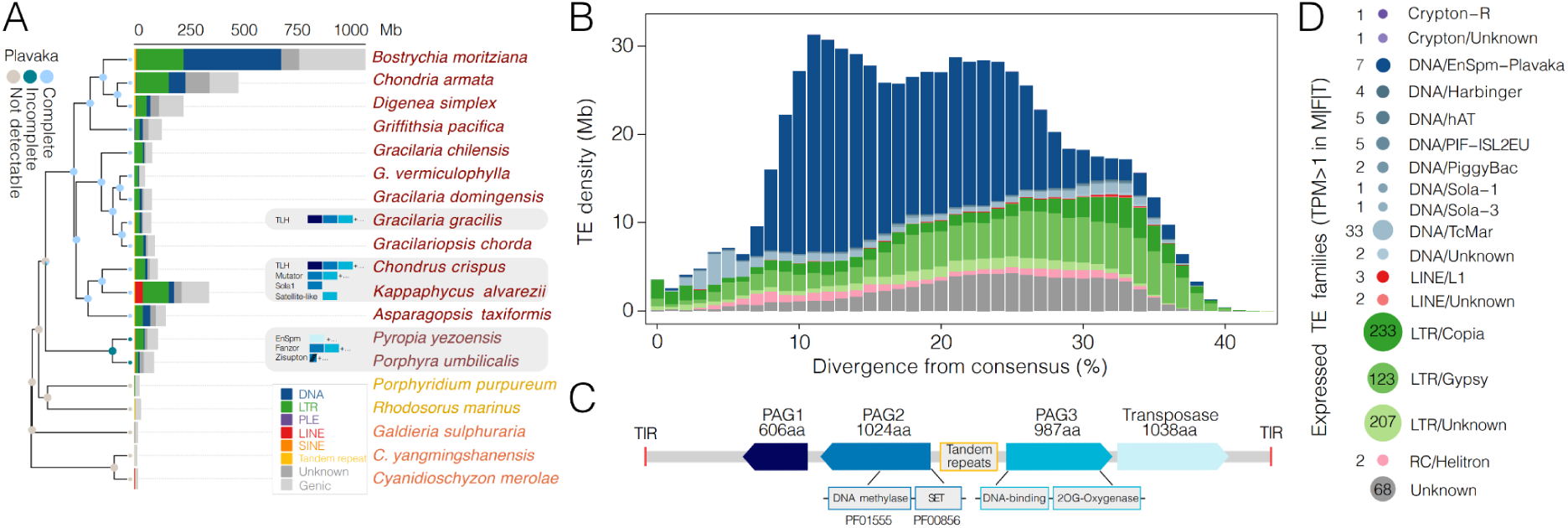
Genome size expansion in *Bostrychia* is primarily driven by the proliferation of *Plavaka* elements. (**A**) Bar chart summarising repeat complement across different species of red algae. The barchart is color-coded according to the relative composition of repeats (in Mb). To the left is a species tree annotated with the results of an ancestral state reconstruction analysis of *Plavaka* elements. The ancestral states are categorized as having a ‘complete’, ‘incomplete’, or ‘not detectable’ *Plavaka* element structure as that seen in *Bostrychia*. (**B**) The sequence divergence landscape of transposable elements (TE) in *Bostrychia.* TE density (in Mb) of each TE grouped by TE class is plotted against the divergence from the consensus sequence of its assigned TE family. Divergence serves as a proxy for the age of the different TE classes. (**C**) A schematic representation of *Plavaka* element structure in *Bostrychia* highlighting the relative position of the three *Plavaka*-associated genes (PAG1-3), the tandem repeats associated with PAG3 and the transposase gene of the *Plavaka-1* family. (**D**) Bubble chart illustrating the number of expressed TE families (TPM > 1 in at least one of male gametophyte, female gametophyte or tetrasporophyte RNA-seq datasets).

In addition to the DDE transposase locus, *Plavaka* elements in *Bostrychia* contain three additional accessory genes, homologs of which have been reported among diverse mobile elements, including giant *Plavaka* and *Mutator* DNA transposons in *Chondrus crispus*^46^ (**Fig. 3C**). While the first *Plavaka* accessory gene (PAG1) has an unknown function, PAG2 encodes a protein with a putative N6-adenine DNA methylase and a SET methyltransferase domain, suggesting that it may modulate transcription via DNA adenine and/or histone lysine methylation activity. The PAG3 protein features an N-terminal domain with homology to ParB DNA-binding proteins, and a putative 2-oxoglutarate (2OG)-dependent oxygenase domain, which perform incredibly diverse biochemical reactions that include modification of DNA, RNA and chromatin^47,48^. ParB proteins bind specific inverted repeat units frequently present in tandem copies on prokaryotic chromosomes and plasmids, which form part of the DNA segregation machinery during cell division^49^. Interestingly, we observed tandem arrays of inverted repeats adjacent to PAG3 genes across *Plavaka* families, suggesting that PAG3 could bind and potentially modify its own transposon sequence via the (2OG)-dependent oxygenase domain. In basidiomycete fungi, just one accessory gene was found associated with *Plavaka* elements, a TET/JBP gene from the class of 2OG-dependent oxygenases involved in cytosine demethylation^45^. Although transposon accessory genes are rare, 2OG-dependent oxygenases and SET domain proteins have been captured by Helitrons, while *HarbingerS* DNA transposons carry an accessory gene encoding a SET domain^50–52^. The only transposons known to bind and modify their own sequence are the *VANDAL* DNA transposons of eudicots, which carry an anti-silencing accessory protein that specifically induces hypomethylation^53^. Our manual curation identified the arrangement of all three accessory genes in the three dominant *Plavaka* families of *Bostrychia* (**Supplemental Fig. 13A**; **Supplemental Table 6**). The majority of *Plavaka* copies were found to be highly degenerate, suggesting that most insertions are likely inactive (**Fig. 3B**; **Supplemental Fig. 13B**). Transcriptomic analysis confirmed this, with low transcript levels observed from seven *Plavaka* families, contrasting with the high expression seen among younger and intact *Copia* and *Gypsy* LTR families (**Fig. 3D**; **Supplemental Fig. 13C**).

While *Plavaka* elements dominate the *Bostrychia* genome, they are less abundant in other red algae, typically comprising less than 3% of repeat content (**Supplemental Fig. 13D**). Our phylogenetic analysis suggests that *Plavaka* elements are only present among the Bangiophyceae and Florideophyceae (**Fig. 3A**). Despite being distributed in both seaweed classes, *Plavaka* elements in the Bangiophyceae occur in the absence of a PAG cassette, with PAG2 and PAG3 instead found associated with *Zisupton* DNA transposons and uncharacterized Fanzor-encoding elements (**Fig. 3A,C**). The capture of the three PAG cassette by *Plavaka* elements thus appears to have coincided with the emergence of the Florideophyceae, suggesting coherent evolution with this dominant class of seaweeds for around 781 million years^2^ (**Fig. 3A**). Interestingly, the *Plavaka* accessory genes exhibit a high degree of mobility, having been incorporated into various TE families across different orders of the Florideophyceae (**Fig. 3A**; **Supplemental Table 6**). These findings highlight the conservation and dynamic nature of *Plavaka* elements across red seaweeds and suggest that their accessory genes have conferred a selective advantage on TE proliferation during red algal genome evolution.

The expansion of TEs thus appears to be the main driver of GSE in *Bostrychia*, with the proliferation of *Plavaka* DNA transposons playing a central role. This is in stark contrast to the modest increases in genome size observed across other red algal orders which, like in plants, is largely attributed to LTR elements (**Fig. 3A**)^5,54–56^. The abundance of DNA transposons in *Bostrychia* is reminiscent of several fish genomes such as zebrafish (*Danio rerio*)^57^, although distinct in that GSE is driven by a single type of DNA transposon. Nevertheless, our analysis suggests that the proliferation of *Plavaka* elements is unlikely to be a general feature of the Ceramiales (**Supplemental Fig. 13D**), although the highly fragmented nature of currently available Ceramiales genomes hinders a definitive conclusion. This raises questions as to why *Plavaka* elements have proliferated so dramatically in *Bostrychia* and more generally about the transcriptional silencing of TEs in red seaweeds. Chromatin organization in red macroalgae still remains unexplored but evidence from the unicellular red alga *C. merolae* shows that TEs are silenced by H3K27me3^58,59^. Future chromatin studies and functional characterization of the *Plavaka* accessory genes hope to provide valuable insights into how these and other TEs have shaped genomic architecture in *Bostrychia* and other red seaweeds.

### Substantial gene family diversification distinguishes the Rhodomelaceae family

The *Bostrychia* genus is classified into the Rhodomelaceae, the largest and most diverse family of red algae, where a sixth of all currently described taxa are found (**Fig. 1B**)^10^. While it is hypothesized that the emergence of distinctive vegetative and reproductive traits in the Rhodomelaceae could have influenced their diversification, the genetic innovations underpinning these traits have remained elusive^11,13^. We thus set out to investigate the evolution of gene families within the Rhodomelaceae and Ceramiales more generally. Interestingly, consistent with the morphological complexity of the Rhodomelaceae, we report 15,338 genes in *Bostrychia*, which represents a one-third expansion in genes when compared with other red seaweeds (**Fig. 2H**). In contrast, 9,641 genes were reported in *Palmaria palmata* (Palmariales) where a significant genomic expansion has also been reported^60^.

Genomic phylostratigraphy revealed that the majority of genes in *Bostrychia* (47.0%; 7,066) have ancient origins that date back to the first cellular organisms and eukaryotes (**Fig. 4A**; **Supplemental Fig. 14A&B**; **Supplemental Table 7**). Just over a third are traceable solely in *Bostrychia* (**Fig. 4A**; 29.3%; 4,430), which may reflect its adaptation to the environmental extremes this species encounters in its native mangrove habitat^21,28^. A further sixth (17.6%; 2,658) represent genes specific to the Rhodophyta and the Rhodymeniophycidae, the largest subclass of the Florideophyceae (**Fig. 4A**). Importantly, we identified 226 genes specific to the Ceramiales, along with a further 732 that originated in the common ancestor of the Rhodomelaceae (**Fig. 4A**), which diversified from 80 and 365 distinct gene families or orthogroups, respectively (**Fig. 4B**). Many of these Ceramiales and Rhodomelaceae-specific orthogroups have no assigned Pfam domain (87.8% and 82.8%, respectively), highlighting substantial *de novo* gene birth during the evolution of this diverse clade (**Supplemental Fig. 15A**). Additionally, our analysis of orthogroup dynamics revealed 106 gene families that expanded in all four of the Ceramiales species we investigated, with a further 124 expanded in the Rhodomelaceae (**Fig. 4C**). The majority of these expanded gene families have deep eukaryotic origins, with only a few (16.0% in the Ceramiales and 15.0% in the Rhodomelaceae) expanded from Rhodophyta-specific gene families (**Fig. 4A**).

**Figure 4.**
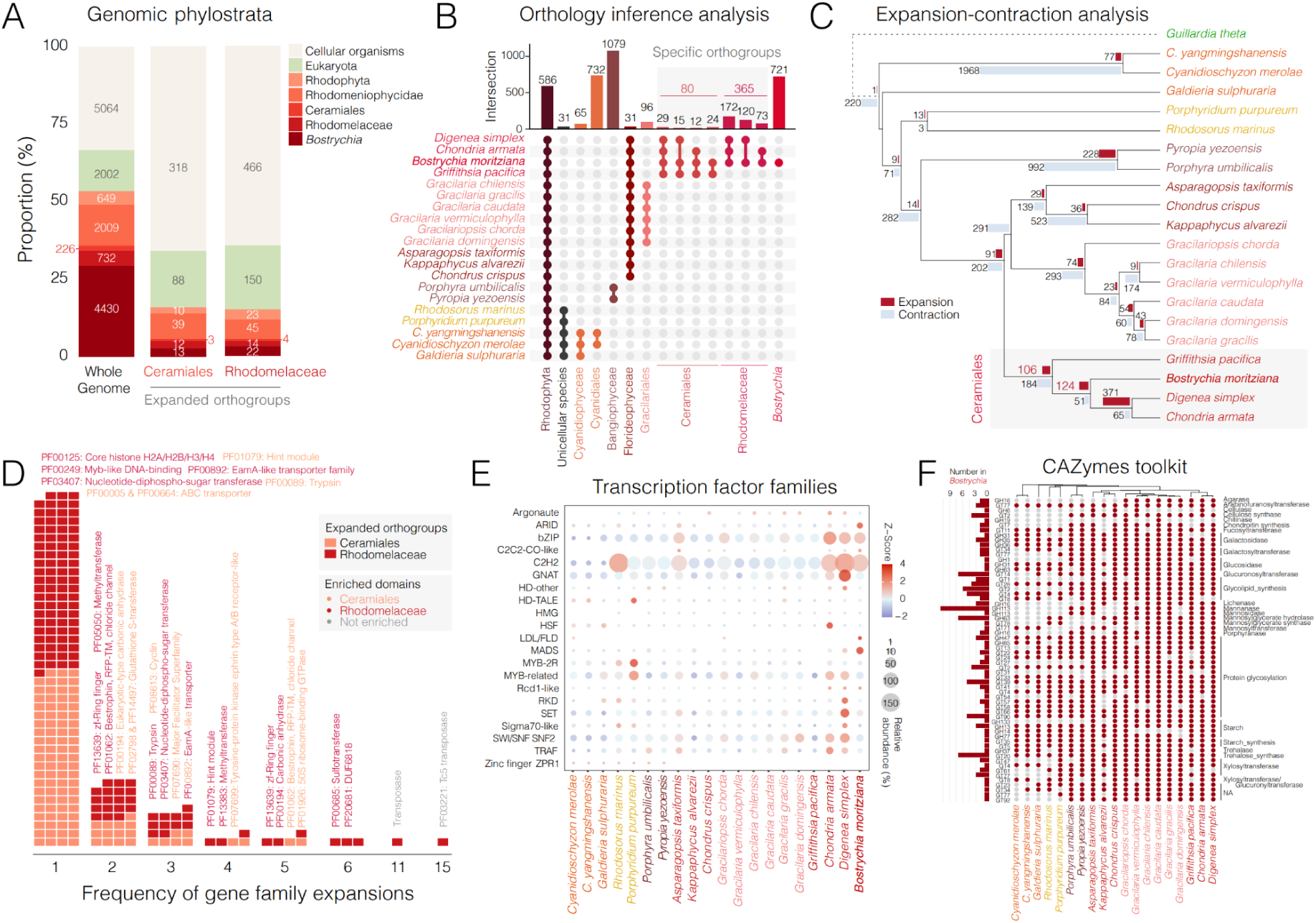
Gene family evolution in the Ceramiales order and Rhodomelaceae family. (**A**) Stacked bar chart showing the proportion of phylogenetically-ranked genes in *Bostrychia* and across the *Bostrychia* genes expanded in the Ceramiales and Rhodomelaceae. (**B**) Upset plot summarising the distribution of orthogroups (OGs) across the red algae. Highlighted are the Ceramiales– and Rhodomelaceae-specific OGs. (**C**) Red algal species tree showing the number of expanded (red) and contracted (light blue) gene families (or OGs) at each node. The Ceramiales node is indicated with grey shading. (**D**) Stacked waffe plot visualizing the frequency of gene family expansions within the Ceramiales and Rhodomelaceae. Examples of significantly enriched Pfam domains within the Ceramiales-expanded OGs (orange) and Rhodomelaceae-expanded OGs (dark red) are indicated. Pfam domains that are not significantly enriched are displayed in grey. (**E**) Bubble chart summarising the number of transcription-associated proteins (TAPs) across the red algae. Bubble sizes show the number of TAPs belonging to a given family. The bubble colors indicate Z-score values, with positive and negative values indicating an above-average or below-average increase in the number of TAPs within a given family, respectively. (**F**) Heatmap summary of the presence and absence of carbohydrate-active enzyme (CAZyme) families across the red algae. CAZyme family names are indicated on the right, while the number of CAZyme genes in *Bostrychia* are represented in the barchart to the left.

To further investigate the function of this distinct gene repertoire, we concentrated on Pfam domains enriched within the four orthogroup categories that were specific to or expanded in both the Ceramiales and Rhodomelaceae (**Supplemental Fig. 15B-E**; **Supplemental Table 7**). Despite most clade-specific orthogroups having an unknown function, our analysis did identify a small proportion with an assigned Pfam domain (12.2% and 17.2% in the Ceramiales and Rhodomelaceae, respectively) (**Supplemental Fig. 15A**). Among the most highly enriched domains within the Ceramiales-specific orthogroups were domains of TRAF zinc finger and bZIP family transcription factors (TFs), as were two types of metalloprotein domains, namely metallopeptidases (type M12 and M23) and multicopper oxidases (**Supplemental Fig. 15B**). Metallopeptidases are a diverse group of peptidases involved in immune response and pathogen defense, suggesting a potential role in bacterial defense in *Bostrychia*^61,62^. The “von Willebrand factor A (vWA) domain” often found in cell adhesion and extracellular matrix glycoproteins^63^ stood out prominently among the orthogroups specific to the Rhodomelaceae (**Supplemental Fig. 15C**), which is found in three tandemly-repeated gene clusters on chromosomes 6, 16 and 25 in *Bostrychia* (**Supplemental Fig. 16A-C**). The most enriched domain among both expanded orthogroups is the Hint module, an autocatalytic domain present in Hint-domain-containing proteins that share similarity to self-splicing inteins (**Fig. 4D**; **Supplemental Fig. 15D&E**)^64^. In animals, this domain occurs in Hedgehog proteins involved in key signal transduction pathways^64^. ABC transporters were also notably expanded among the Ceramiales, which are known to play a role in plant growth and development by transporting various substrates including auxin and other phytohormones^65^. Highly enriched within the Rhodomelaceae-expanded orthogroups were domains of core histones as well as several transcription-associated proteins (TAPs), including Ring-type zinc finger and MYB-related domains (**Supplemental Fig. 15E**). ARID, SWI/SNF-SNF2 and other zinc-finger domains like C2H2 were also overrepresented. A broad assessment of TAPs across red algae further highlights the expansion of specific TAP families, particularly C2H2, bZIP and Rcd1-like TFs within the Rhodomelaceae, with MADS box TFs further prominent in *Bostrychia* (**Fig. 4E**). Interestingly, the expansion of C2H2, bZIP and MADS box TFs is proposed to have been key for increased complexity in land plants, alluding to their expansion in the Rhodomelaceae as having a similar impact^66^. Our results thus highlight substantial gene family diversification in both the Ceramiales and Rhodomelaceae, which is reminiscent of the genetic adaptations associated with developmental complexity in their sister land plant lineage.

### *Bostrychia* encodes an expanded carbohydrate metabolism toolkit

Carbohydrate metabolism is fundamental to the growth and development of all plants and algae. Carbohydrate-active enzymes (CAZymes) play a central role in regulating protein glycosylation, glycolipids, the cell wall and energy metabolism. Glycosyltransferases (GTs) catalyze the formation of glycosidic linkages between sugars and substrates, while glycoside hydrolases (GHs) cleave glycosidic bonds. *Bostrychia* contains a diversity of 96 GTs and 48 GHs, distributed across 32 and 16 CAZyme families, respectively (**Fig. 4F**; **Supplemental Table 8**).

*Bostrychia* thrives in unique habitats characterized by harsh environmental extremes, including rapid salinity fluctuations and desiccation, which requires finely-tuned metabolic pathways to survive osmotic stress. Evidence of environmental adaptation is marked by CAZymes involved in regulating low molecular weight carbohydrates (LMWCs), which serve as compatible solutes for osmotic balance and as energy reserves. Notably, *Bostrychia* contains seven genes (GT20) for synthesizing the LMWC trehalose, along with a trehalase (GH37) for its degradation, despite findings that trehalose is largely absent in the Ceramiales and undetected in Rhodomelaceae^67^. Functional studies of putative trehalose synthases in other red algae suggest that these enzymes may instead synthesize other LMWCs, such as floridoside and isofloridoside^68,69^. Further functional studies may reveal intriguing insights into the role of these LMWC-related enzymes and their contribution to the adaptation of *Bostrychia* to environmental fluctuations.

For energy metabolism, *Bostrychia* possesses eight CAZymes necessary for starch synthesis and degradation^4,5^. Mannosylglycerate, an important LMWC, primarily functions as an energy reserve in other species in the *Bostrychia* genus^70–72^. In *Bostrychia*, this is supported by a mannosylglycerate synthase (GT78) and a unique cluster of seven mannosylglycerate hydrolases (GH63) to facilitate its degradation (**Supplemental Fig. 16D&17A**). Mannosylglycerate hydrolase clusters have not been reported in other red macroalgae, raising questions about the evolutionary origin and regulation of this cluster (**Fig. 4F**).

Red macroalgae can possess cell walls with cellulose, xylan or mannan as crystalline or paracrystalline polysaccharides, which help provide strength and structure to tissue^73^. *Bostrychia* contains four putative *CELLULOSE SYNTHASE (CESA)* genes, two of which share strong identity with bacterial *CESAs*, while the other two share identity with eukaryotic *CESAs*, including the functionally characterized *CESA1* from *Calliarthron tuberculosum* (Florideophyceae)^74^. Although *Bostrychia* lacks cell wall mannans^75^, it shows a striking enrichment of β-1,4-mannanases (GH113), with 11 genes identified across seven chromosomes (**Supplemental Fig. 17B**). As mannanases are typically associated with cell wall mannan degradation, this abundance may indicate a role in epiphytism, either in defending against macroalgae competitors or by facilitating attachment to other macroalgae. This putative dual functionality of β-1,4-mannanases highlights a potential adaptive advantage in environments where competition for space and resources is intense, allowing *Bostrychia* to interact with or defend itself from surrounding macroalgal species.

### Gene-rich UV sex chromosomes harbor orthologs of ancient life cycle regulators

Like bryophytes and most green and brown seaweeds, sex determination in red seaweeds occurs during the haploid phase of the life cycle by the inheritance of a U or V sex chromosome in the female and male gametophyte, respectively^76,77^. U and V sex chromosomes are often composed of a region with shared homology called a pseudo-autosomal region (PAR) and a non-recombining sex-determining region (SDR) that harbors key genes that specify either sex (Umen & Coelho, 2019). In red algae, a UV system was originally suggested by Mendelian segregation in *Gracilaria tikvahie*^78^, with evidence for sex-linked loci since shown in several other taxa, including other *Gracilaria* species^79–81^, *Aglaothamnion oosumiense*^35^, *Pyropia*^82^ as well as *Bostrychia*^26^. More recently, genomic analyses have resolved the SDRs in four *Gracilaria* species, which are relatively small but harbor a handful of conserved genes^83^. Like the brown alga *Ectocarpus siliculosus,* the SDR of the UV sex chromosomes in *Gracilaria* and *Pyropia* is sandwiched between two PARs^82–85^, whereas in the bryophytes *Marchantia polymorpha* and *Ceratodon purpureus*, which harbour the oldest known sex chromosomes, they are only composed of a highly expanded SDR^76,86^.

The UV sex chromosomes are among the smallest chromosomes in *Bostrychia*, measuring out at 19.98 Mb and 20.87 Mb in the female and male, respectively (**Fig. 5A**; **Supplemental Fig. 4**). Synteny and k-mer-based analyses revealed the first portion to be highly syntenic and indicative of a PAR, followed by extended sex-specific regions representing the female (12.37 Mb) and male SDR (13.13 Mb) (**Fig. 5A**; **Supplemental Fig. 18A&B**). GC content of the PAR was slightly lower compared to autosomes but significantly higher in the SDR of both sexes, which was further distinguished by a significantly higher density of TE insertions, particularly helitron elements (**Fig. 5A**; **Supplemental Fig. 18C-G**). With 195 and 187 genes, respectively, the U and V chromosomes have the lowest gene density of all chromosomes in *Bostrychia* (**Supplemental Fig. 8**). Although these genes were significantly younger than the average observed across all chromosomes, similar trends were observed on autosomes 11, 20 and 31, suggesting that the enrichment of young genes is not a UV-specific feature in *Bostrychia* as reported in brown seaweeds^87^ (**Supplemental Fig. 14A&B**). The presence, order and synonymous distance (dS) between homologs on the PAR are suggestive of ongoing recombination (**Fig. 5A**), although a lack of recombination maps prevents further conclusions. Interestingly, we noted 22 PAR genes that were specific to either the U or V (**Fig. 5A**), which is unlikely due to assembly errors since we were able to recover a gapless PAR on the V chromosome (**Fig. 5A**). Nevertheless, these sex-specific PAR genes were largely un-transcribed (86.36% in the male and 72.73% in the female) and mostly had no homology in other species (63.64%) (**Supplemental Table 9**). Around a quarter of PAR genes were transcribed in at least one of the gametophytes or tetrasporophyte (28/87 and 22/85 in the female and male, respectively; 32.18% and 25.88%), with expression observed primarily from the borders of the PAR, whereas the center was mostly untranscribed (**Fig. 5A**). We further confirmed sex-specific expression and conservation of UV genes using previous data from a geographically-distinct strain of *Bostrychia* (**Supplemental Table 9**)^26^.

**Figure 5.**
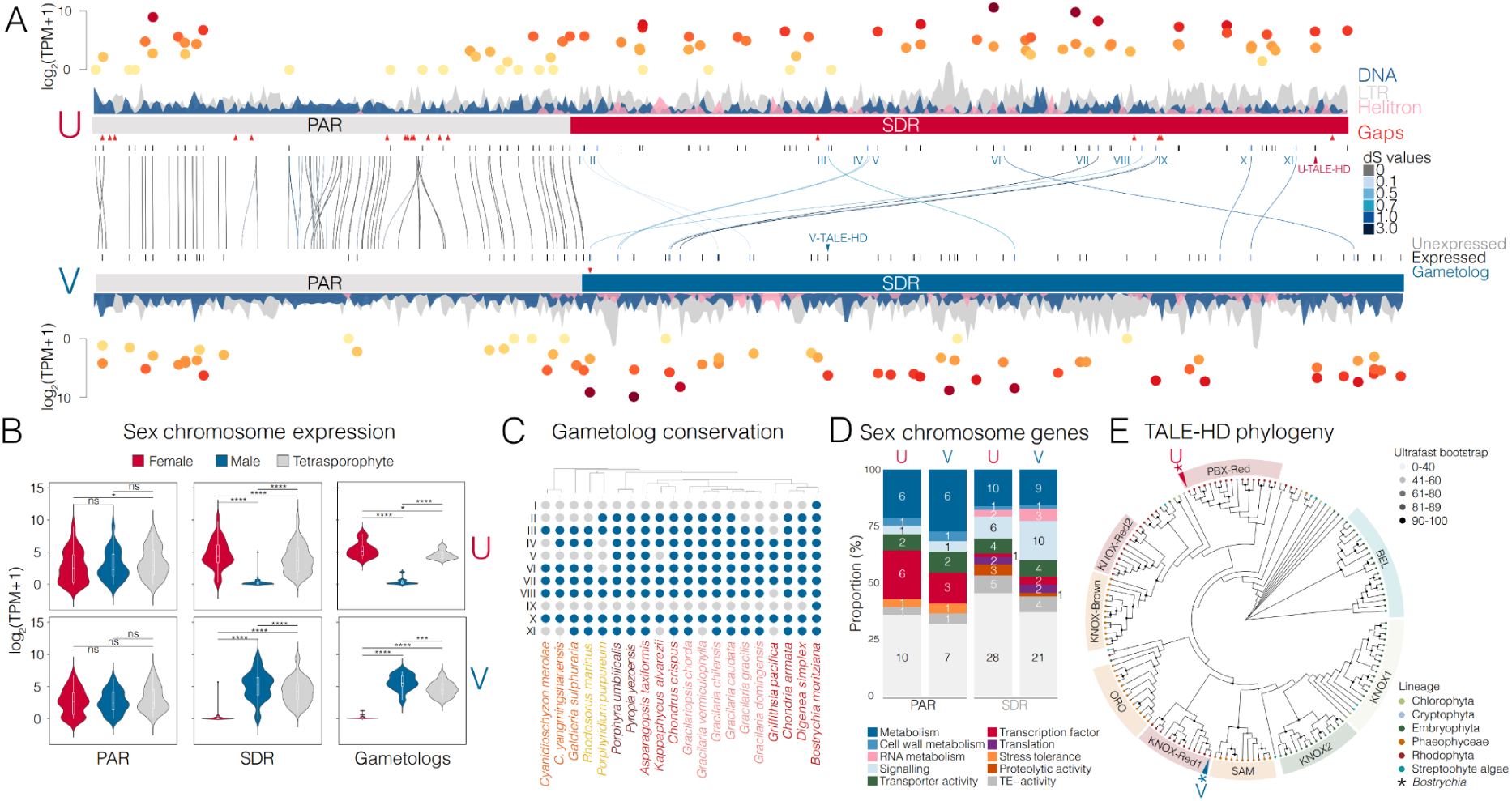
The UV sex chromosomes of *Bostrychia moritziana*. (**A**) Karyoplot comparing synteny between the female U and male V sex chromosomes. The PAR regions are indicated with a grey bar, while the female SDR is highlighted in red and the male SDR in blue. Genes are represented by short lines and colour coded based on whether they are unexpressed (grey), expressed (black) or whether they represent a gametolog (blue). Gametologs are further labeled as I-XI, while the two TALE-HD genes are labeled separately. Gene synteny is indicated with connecting lines and is color-coded based on synonymous distance (dS) values. Gaps are indicated with red triangles. The density of DNA transposons (blue), LTRs (grey) and Helitrons (pink) is shown as a density plot above (U) or below (V) the chromosomes. Gene expression levels are visualized above (U) or below (V) using a color scale that reflects mean log2-transformed TPM+1 values. (**B**) Violin plots showing the expression by log2-transformed TPM+1 values of PAR genes, SDR genes and gametologs on the U and V sex chromosomes in the female gametophyte (red), male gametophyte (blue) and tetrasporophyte (grey). Wilcoxon test results for statistical comparisons are shown above the violin plots (* (*p*-value < 0.05), ** (*p*-value < 0.01), *** (*p*-value < 0.001), ns = no significance). (**C**) Heatmap summary of the presence and absence plot of gametolog gene conservation across the red algae. (**D**) Stacked bar charts summarising the broad functional categories of expressed genes in the PAR and SDR of the U and V sex chromosomes. (**E**) Rooted maximum-likelihood tree of TALE-HD TFs in brown algae, cryptophytes, red algae, green algae, streptophyte algae, and land plants. Dots at the tips indicate the respective phyla and are colour-coded according to the key. Outer shading around the circumference highlights the different classes of TALE-HD TFs with the position of the sex-linked pair in *Bostrychia* also indicated. Ultrafast bootstrap values are indicated as color-coded dots at each node of the phylogeny.

Consistent with their sex-specificity, we observed very poor synteny between the SDRs aside from 11 orthologous genes, one of which was duplicated in the male (**Fig. 5C**). The majority of these so-called gametologs (8 of 11) appear to be ancient and highly conserved across red algae and are associated with various biological functions, including metabolism, cell-wall metabolism, RNA metabolism, signalling and transporter activity (**Fig. 5C**). One gametolog emerged specifically in the Rhodophyta and encodes for a Pleckstrin domain-containing protein, while the other two encode unknown proteins specific to *Bostrychia* (**Fig. 5C**). Sequence divergence between gametologs was significantly elevated compared to PAR genes (**Supplemental Fig. 18H**), highlighting both recent and ancient events of gene migration into sex-linked regions. In addition to the 11 gametologs, the SDRs each contained 97 U– and 90 V-specific genes (**Fig. 5A**), although only around a half were appreciably expressed in the gametophyte and/or tetrasporophyte (51/97 on the U; 45/90 on the V). Both SDR and gametolog genes were only expressed in each respective gametophyte, further validating their sex-specifity (**Fig. 5B**). Overall, SDR genes had higher expression compared to PAR genes (**Supplemental Fig. 18I**). Unlike PAR genes, which had similar expression levels in the gametophyte and tetrasporophyte generations, gametolog and SDR genes were significantly down-regulated in the tetrasporophyte (**Fig. 5B**), although this may be conflated by the presence of maturing haploid tetraspores on the tetrasporophyte we processed. The UV sex chromosomes in *Bostrychia* thus share features with the UV systems reported in other red algae, brown algae and land plants, particularly with regard to the presence of a PAR and the differential expression of sex-linked genes across the haploid and diploid generations of the life cycle^83,86,88,89^. This may relate to the fact that *Bostrychia* has an isomorphic life history on the one hand, yet differentiates distinctive reproductive structures between sexes and generations. The presence of two sporophytic phases is likely to add an additional layer of generation-antagonistic selection between the U and V chromosomes, which could offset sexually-antagonistic selection and influence the evolution of UV systems in this order of complex red seaweeds.

Next, we focused on the function of sex chromosome genes in *Bostrychia* since these are likely to play a key role in sexual differentiation and life cycle control. Due to the relatively small sample size of genes, GO term enrichment analysis did not yield significant or meaningful results. We thus assessed putative gene functions on the sex chromosomes by manually assigning the transcribed set of UV genes into broad functional categories (**Fig. 5D**). Interestingly, the relative proportion of these functional categories differed between the PAR and SDR. For example, the SDRs encoded more genes involved in signalling processes, with the male SDR having almost twice that of the female (**Fig. 5D**; **Supplemental Table 9**). Similarly, the PARs contained more genes encoding TFs than the SDRs, with virtually all classified as bZIP TFs (**Fig. 5D**; **Supplemental Table 9**). However, most notable were a distinct pair of <u>T</u>hree <u>A</u>mino acid <u>L</u>oop <u>E</u>xtension <u>H</u>omeo<u>d</u>omain (TALE-HD) TFs that were present on the female and male SDR (**Fig. 5A**). TALE-HDs are highly conserved across eukaryotes and are classified as atypical homeodomains due to a characteristic three-residue insertion^90–92^. In land plants, TALE-HD TFs are categorized into two classes, KNOTTED-like (KNOX) and BEL1-like (BEL), based on the presence of additional associated domains^92,93^. The sex-linked TALE-HD TFs were the only two present in *Bostrychia*, with the number of TALE-HD genes varying between one and nine across unicellular and multicellular red algae, as reported previously^94,95^ (**Fig. 5E**). TALE-HDs in red algae have been categorized into two classes (1) KNOX-Red, which shares homology with plant KNOX genes and is further subdivided into KNOX-Red1 and KNOX-Red2, and (2) PBX-Red, which is homologous to PBC-domain-containing TALE-HD TFs in animals^96^. KNOX-Red1 is distinguished by an ELK domain, KNOX-Red2 contains KN-A and KN-C2 domains and PBX-Red features a PBL (PBC-like) domain (**Supplemental Fig. 19**)^96^. We found KNOX-Red1 in most red algal classes, while KNOX-Red2 was restricted to the Cyanidiophyceae and Florideophyceae (**Fig. 5E**), as reported previously^96^. In *Bostrychia,* the female-specific TALE-HD TF was classified as PBX-type, while the male-specific TF was classified as KNOX-Red1 (**Fig. 5E**).

Although KNOX and BEL regulate organ and reproductive development in flowering plants^91,97^, they have deep evolutionary origins in eukaryotes^96,98–100^. In the green lineage, KNOX and BEL form a heterodimer that activates the diploid zygotic program in the chlorophyte microalga *Chlamydomonas reinhardtii* and the bryophyte *Marchantia polymorpha*^101–103^. Similarly, the TALE-HD TFs OUROBOROS and SAMSARA also heterodimerise in the brown alga *Ectocarpus* to regulate the gametophyte-to-sporophyte transition^89,104,105^, with metazoan TALE-HDs also capable of heterodimerisation^96,106^. Interestingly, two TALE-HD TFs in the red microalga *Galdieria* and one in the bangiophyte macroalga *Pyropia* are differentially expressed between haploid and diploid phases of the life cycle, with those in *Galdieria* shown to play an essential role in the haploid-to-diploid transition^94,95^. Both of the sex-linked TALE-HD TFs in *Bostrychia* are expressed in the tetrasporophyte generation (**Supplemental Table 9**), raising the intriguing possibility that they may also heterodimerise to initiate sporophytic development, prompting further questions about how this activity would be modulated between the two sporophytic stages that typify the triphasic life cycle of the Florideophyceae.

## Conclusions

In summary, the *Bostrychia* genome has revealed key genetic adaptations that distinguishes the morphologically complex Rhodomelaceae and confirms classical observations that genomic expansion may be a common feature of red algal evolution^16^. The adaptations specific to the Rhodomelaceae and Ceramiales have likely conferred a competitive advantage that has contributed to the evolutionary success and dominance of these groups of red algae, in turn highlighting *Bostrychia* as a powerful model system to explore red algal development, reproduction and evolution. The red algal transition to multicellularity ranks among the earliest known across eukaryotes^2,107^, which occurred in spite of the loss of deeply conserved pathways normally deemed essential in other multicellular lineages^8^. In light of this and the common evolutionary origin of red seaweeds and plants, our findings offer new perspectives and valuable genomic resources for elucidating the universal origins of complex multicellularity in these distinct yet closely related lineages of eukaryotes.

## Methods

### Culture and sample preparation

Male and female sibling gametophytes of *Bostrychia moritziana* strain 3235 (isolated from Beachwood, Durban, Natal, South Africa) were described previously^30,31^. The tetrasporophyte was isolated from a single carpospore after crossing the male and female gametophyte. Algal culture was performed in autoclaved natural seawater (NSW) enriched with half-strength Provasoli enriched seawater (PES) medium^107^ and cultivated in 2-liter flasks with aeration. Fresh enriched seawater was changed weekly to promote growth. Growth conditions were maintained at 20°C with a light intensity of <20 µM in a 16h light and 8h dark cycle. Bulked gametophyte tissue for genomic DNA extraction was treated with 200 mg/l Ampicillin, 100 mg/l Kanamycin, 500 mg/l Penicillin and 100 mg/l Streptomycin for three days prior to harvest and storage at –20 °C.

### Genomic DNA extraction and sequencing

The extraction of high-molecular weight genomic DNA (average size = ∼70 kb) was accomplished using a tailored protocol. First, algal tissue was ground in liquid nitrogen, then pre-washed twice with sorbitol wash buffer (0.35 M D-Sorbitol, 100 mM Tris-HCl pH 8, 20 mM EDTA pH 8, 2% PVP-40 and 1% 2-Mercaptoethanol) to remove phenolic compounds, followed by centrifugation at 5,000 x g for 5 minutes. Cell lysis was accomplished by adding CTAB lysis buffer (4% CTAB, 100 mM Tris-HCl pH 8, 20 mM EDTA pH 8, 1.4 M NaCl and 2% PVP-40), 100 µg/ml RNAse A and 100 µg/ml Proteinase K to the pre-washed pellet for 1 hour at 60°C. An equal volume of chloroform:isoamyl alcohol (24:1, v/v) was then added and mixed gently prior to phase separation with centrifugation at 10,000 x g for 10 minutes. The upper aqueous phase was treated with 100 µg/ml RNAse A at 60°C for 15 min. The chloroform clean-up step was then repeated without RNAse treatment. Genomic DNA was then precipitated by the addition of 0.7x ice-cold isopropanol and pelleted by centrifugation at 5,500 x g for 30 minutes at 4°C, followed by washing with 70% ethanol. To remove agar-derived contaminants, resuspended DNA was further cleaned using a Zymoclean Large Fragment DNA Recovery Kit (No. D4045) followed by a further clean-up with AMPure XP beads. The resulting HMW DNA was subjected to sequencing using two long-read technologies, namely Oxford Nanopore Technologies (ONT) MinION and PacBio SMRTbell sequencing. One male and one female ONT library were prepared using the Ligation sequencing gDNA (SQK-LSK110-XL) library preparation kit for FLO-MIN106D flow cells. An additional female ONT library was prepared using the SQK-LSK114 library preparation kit for a FLO-MIN114 flow cell. Sequencing was performed on a ONT MinION Mk1B device (**Supplemental Table 3**). Basecalling was performed using Guppy v6.5.7 with the *trim_adapters*, *trim_primers* and *calib_detect* options^108^. For the two libraries sequenced on FLO-MIN106D flow cells, the dna_r9.4.1_450bps_hac configuration file was used. For two runs of the same library using the FLO-MIN114 flow cell, the dna_r10.4.1_e8.2_400bps_sup and the dna_r10.4.1_e8.2_260bps_sup configuration files were applied. PacBio libraries for both the male and female gametophyte HMW DNA were prepared using the SMRTbell prep kit 3.0. Sequencing was performed on a Pacific Bioscience Sequel II platform (**Supplemental Table 3**). PacBio subreads were processed into HiFi reads using ccs 6.4.0 with the *--min-rq=0.88* option^109^. Demultiplexing of reads was performed using lima 2.7.1 with the *--split-named* option^110^. Subreads were realigned to the generated HiFi reads separately for the male and female datasets using actc v0.2.0^111^. Finally, error correction of the HiFi reads was performed using DeepConsensus v1.2.0 with the *--batch_size=2048* and *--batch_zmws=1000* options^112^.

### RNA-seq analysis

Total RNA of the male and female gametophyte was extracted using a Qiagen RNeasy PowerPlant Kit (No. 13500-50) according to manufacturer instructions. To concentrate RNA, 155 mM RNase-free sodium acetate, 0.75X isopropanol, and 116 µg/ml Glycoblue were added, the solution was mixed thoroughly and incubated at –80°C for 60 minutes. The sample was centrifuged at maximum speed (13,000 g) for 30 minutes at 4°C. The supernatant was carefully removed, and the pellet was washed with 500 µl of ice-cold 75% ethanol. The sample was centrifuged again at maximum speed for 15 minutes at 4°C. The pellet was air-dried on ice for 10 minutes and resuspended in ice-cold RNase-free water. Libraries were prepared with the NEBNext^®^ Ultra™ II Directional RNA Library Prep Kit (New England Biolabs, E7760S). Sequencing of 150 bp paired-end reads was done on an Illumina NextSeq 2000 platform (**Supplemental Fig. 6A**). Adapters of RNA-seq reads of *Bostrychia, Kappaphycus* and *Asparagopsis* were trimmed using TrimGalore v0.6.10 with the *--fastqc_args* option^113^.

### Hi-C library preparation and sequencing

Hi-C libraries were generated as described previously with some modifications^114,115^. Female and male gametophyte tissue was cross-linked in 2% formaldehyde for 30 minutes at room temperature and then quenched with 400 mM glycine. Tissue was then ground in ice-cold nuclei isolation buffer (0.1% triton X-100, 125 mM sorbitol, 20 mM potassium citrate, 30 mM MgCl2, 5 mM EDTA, 5 mM 2-mercaptoethanol, 55 mM HEPES pH 7.5). The nuclei suspension was then incubated with 0.5% SDS at 65°C for 5 minutes, followed by quenching with 10% Triton X-100 and incubation at 37°C for 15 minutes. DpnII digestion was carried out overnight at 37°C using 10x DpnII buffer and 100U of DpnII and then deactivated by incubation at 62°C for 20 minutes. Biotinylation was performed by the addition of 0.4 mM biotin-14-dCTP, 10 mM dTTP, dGTP and dATP and 40U Klenow, followed by incubation at 37°C for 2 hours. The ligation was performed by adding 10X ligation buffer (300 mM Tris-HCl pH 7.8, 100 mM MgCl_2_, 100 mM DTT and 1 mM ATP), 10% Triton X-100 and 5U of the T4 DNA ligase, then incubated at room temperature for 4 hours. The biotinylated nuclei were then treated with SDS buffer (50 mM Tris-HCl pH 8.0, 1 mM EDTA and 1% SDS) and proteinase K at 55°C for 30 min, followed by incubation with 0.2 M NaCl at 65°C overnight to reverse crosslinks. DNA precipitation was performed using 1x volume of Phenol:Chloroform IAA, 0.3 M NaAC and 1x volume iso-propanol, with subsequent washing using 70% ethanol. Shearing of DNA and size selection were performed using a Covaris E220 ultrasonicator and AMPure XP beads, respectively. Library preparation was performed using a NEBNext^®^ Ultra™ II DNA Library Prep Kit (NEB, no. E7645) according to manufacturer instructions. 150 bp paired-end reads were generated using an Illumina NextSeq 2000 platform.

### *De novo* genome assembly and scaffolding

A draft hybrid assembly from ONT and PacBio HiFi reads was generated separately from the male and female gametophyte sequencing data (**Supplemental Table 3**). A number of different assemblers were benchmarked to find the best performing assembly tool for the male and female data set. Performance was evaluated based on the completeness of the expected gene content and the contiguity of the genome using BUSCO v5.8.2 and N50 scores, respectively^116^. For the male dataset, a 1.2 Gb assembly with 3,098 contigs (N50 = 1.0 Mb, complete BUSCO percentage = 78.9%) was generated with Hifiasm v0.16.1-r375 (using the *-l0* and the *--hg-size* option) and polishing with Racon v1.5.0^117,118^. Based on the overrepresentation of Nanopore reads compared to the male dataset, the assembler Flye v2.9.2-b1795 (using the *--scaffold* option) together with Racon v1.5.0 was used for the female, resulting in a 1.17 Gb assembly with 11,876 contigs (N50 = 1.1 Mb, complete BUSCO percentage = 78.4%)^119^. To resolve the assemblies at a chromosome-level and to filter contigs belonging to the associated microbiome, a 3D *de novo* assembly Hi-C pipeline was applied using juicer and 3D-DNA^120,121^. We first identified the genomic localization of DpnII restriction sites using the *generate_site_positions.py* script from the Juicer tool. Subsequently, we employed Juicer to generate Hi-C contact maps^121^.

3D-DNA assembly was performed using the *run-asm-pipeline.sh* script with the *merged_nodups.txt* file generated by juicer^120^. This step facilitated manual correction of scaffolding and refinement of the order and orientation of each assembly. The Hi-C contact maps were visualized using Juicebox v1.11.08^122^. A few small inversions in the male assembly that were caused by assembly gaps were corrected using contiguous regions in the female assembly as reference (i.e. on chromosomes 3, 5, 9, 10, 14, 15, 28 and 29). In addition, selected chromosomes in the male assembly were reversed to ensure a unified orientation with those in the female assembly (i.e. chromosomes 1, 7, 9, 10, 13, 18, 19, 21, 24, 27, 30 and 31). We used the *run-asm-pipeline-post-review.sh* script in the 3D-DNA package to incorporate manual edits and finalize the genome assembly in fasta format. Finally, gap closing was performed using TGS-GapCloser v1.2.1, while errors were corrected with Racon v1.5.0 using both Nanopore and HiFi sequencing reads^123^.

We standardized chromosome nomenclature of both assemblies using a uniform format (i.e., Chr[number]). Subsequent analyses revealed the integration of a bacterial contig on the female sex chromosome caused by assembly gaps, which was manually removed to ensure assembly accuracy. Due to the higher contiguity and completeness of the male assembly, as indicated by a higher N50 and fewer gaps, we used it as the reference genome for our analyses. To ensure the representation of both sex-specific regions, we added the female SDR to the male assembly (**Table 1**). Synteny plots between the male and the female assembly were generated using GENESPACE v1.3.1^124^. The complete mitochondrial genome was found to be integrated into chromosome 10 in both the male and female assembly and was also removed prior to further analyses. The mitochondrial genome in the male assembly has a total length of 26,150 bp. The plastid genome was included in the unplaced contigs of *Bostrychia* and was identified using Tiara v1.0.3^125^. The complete plastid genome in the male assembly has a total length of 188,388 bp.

### Structural, functional and repeat annotation in *Bostrychia*

Prior to annotating protein-coding genes in the *Bostrychia* genome, each assembly was initially soft-masked for repetitive regions and TEs. First, a comprehensive TE library was generated containing consensus sequences of all TE families and repetitive sequences. This was performed with RepeatModeler v2.0.4 using the *LTRStruct* option to include structural LTR detection^126^. Repeats and TEs were then soft-maked with RepeatMasker version open-4.0.9 using the *xsmall* and *gff* options^127^. Tandemly repeated satellite and microsatellite elements were annotated with Tandem Repeats Finder (TRF) v4.09.1^128^ using the following parameters: 2 (*Match*), 7 (*Mismatch*), 7 (*Delta*), 80 (*PM*), 10 (*PI*), 50 (*Minscore*), 2000 (*MaxPeriod*), *m, f, d* and *ngs*. To ensure soft-masking of the tandem repeats across the genomes, bedtools v2.31.1 using the *maskfasta* option was applied^129^.

Gene prediction in *Bostrychia* was performed using RNA-seq data alone (BRAKER1) and using both RNA-seq data and a published manually curated orthologous set of red algal protein sequences (BRAKER3)^130–142^. First, trimmed RNA-seq reads of the male gametophyte, female gametophyte and the tetrasporophyte were mapped to each assembly with Hisat2 v2.2.1 using the *dta* option providing strandness information (RF)^143^. The first round of gene predictions was performed using only RNA-seq evidence with the *softmasking* parameter, resulting in 23,496 male and 23,028 female genes. The second round of annotation used in addition also a conserved orthologous set of protein sequences derived from 16 published red algal species (**Supplemental Table 3 DS1**). For this, orthologous genes and orthogroups were identified using OrthoFinder v2.5.5^144^. Orthogroups containing at least 80% of the species were then filtered using OGFilter^145^, yielding a total of 2,712 orthogroups. These orthogroups were combined into a single fasta file, then combined with RNA-seq evidence for the second round of annotation, resulting in 8,231 male and 8,170 female gene predictions. Both rounds of gene prediction were then integrated using TSEBRA, resulting in 17,712 genes in the male and 17,334 genes in the female^146^. Gene IDs were standardized using a custom script to follow a systematic naming convention. The format is structured as Bm01g000010.t1, where Bm refers to the species (*Bostrychia moritziana*), 01 indicates the chromosome number, g denotes “gene”, 000010 represents the unique gene identifier, and .t1 specifies the transcript number. To simplify downstream analyses, only the longest transcript isoform was used. The gene set was additionally filtered to exclude TE-related genes, which were defined as sequences that were 100% soft-masked by the repeat annotation (894 and 629 in the male and female, respectively). The final gene annotations resulted in 15,230 and 15,031 genes in the male and female assembly, respectively.

Functional and protein domain annotation was conducted using InterProScan v5.64-96.0 and EggNOG-mapper v2.1.8^147,148^. A curated annotation of transcription-associated proteins (TAPs) was carried out using TAPScan v4, while genome-wide annotation of tRNA and rRNA loci was performed using tRNAscan-SE v2.0.12 and Barrnap v0.9, respectively^149–151^. Gene prediction, functional annotation, as well as rRNA and tRNA annotation of the plastid genome was performed using GeSeq^152^. For tRNA annotation, ARAGORN v1.2.38 and tRNAscan-SE v2.0.7 were used as third-party annotators, while Chloë v0.1.0 was enabled as an additional stand-alone annotator. The same annotations, including gene prediction, functional annotation, and rRNA and tRNA annotation, were performed for the mitochondrial genome using MFannot^153^.

### Structural, functional and repeat annotation of published red algal genomes

For subsequent comparative analyses, various published red algal genome datasets were used (**Supplemental Table 3**). However, protein-coding gene annotation was missing for some selected genomes, namely for *Kappaphycus alvarezii*^154^*, Asparagopsis taxiformis*^155^*, Chondria armata*^156^*, Digenea simplex*^157^ and *Griffithsia pacifica*^158^. Repeats were first soft-masked as described above for *Bostrychia*. For gene prediction in *Kappaphycus* and *Asparagopsis*, only a single round of annotation was performed using both RNA-seq evidence and the orthologous set of red algal protein sequences described above. The resulting gene annotations were then filtered to exclude TE-genes and to retain only the longest transcript isoforms, resulting in 6,672 genes and 14,903 genes in *Kappaphycus* and *Asparagopsis*, respectively.

For *Chondria* and *Digenea*, an additional clean-up step was performed prior to gene prediction to address the high levels of contamination in these genome assemblies. Blobplots were generated using BlobTools v1.1.1 to visualize the taxonomic composition of contigs and guide the filtering of potential contaminants from the assemblies^159^ (**Supplemental Fig. 20A&B**). For this, genomic reads were aligned to their respective genome assembly to assess coverage using Minimap2 v2.28-r1209 with the *map-ont* option^160^. Taxonomic assignments were then performed by conducting sequence similarity searches of the assemblies against the NCBI nt database with BLAST using the parameters recommended in the BlobTools documentation (-*evalue* 1e-25 –*outfmt* ‘6 qseqid staxids bitscore std’ –*max_target_seqs* 1 –*max_hsps* 1)^161,162^. The final blobplots were generated using the *create, view* and *plot* functions integrated in BlobTools^159^. Based on manual inspection of the taxonomic distribution and classification in the blobplots, contigs were filtered as follows. For *Chondria*, all contigs with a GC content greater than 0.55 and those assigned to *Pseudomonadota* were removed. For *Digenea*, contigs with a GC content exceeding 0.5 and those assigned to *Pseudomonadota* were excluded. Because RNA-seq data was not available for either species, gene prediction was performed using the orthologous set of red algal protein sequences described earlier. TE-related genes were then filtered out and only the longest transcript isoforms retained, resulting in 20,631 genes in *Chondria* and 28,013 genes in *Digenea*.

To improve the highly fragmented contigs of the *Griffithsia* genome assembly, a scaffolding step was performed using RNA-seq reads and P_RNA_scaffolder with default settings^163^. Following scaffolding, contaminant contigs were filtered out as described for *Chondria* and *Digenea*, using the same parameters applied to *Chondria* (**Supplemental Fig. 20C**). The mapping of Illumina reads was performed using hisat2 v2.2.1 using the *dta* option instead of minimap2^143^. Gene predictions were performed as described for *Kappaphycus* and *Asparagopsis,* resulting in 9,334 genes in *Griffithsia*. Functional and protein domain annotation was conducted for all 19 red algal protein sets using InterProScan v5.64-96.0 and TAPscan v4^147,150^.

### Analysis of repeats and TEs

To ensure consistent repeat annotation across the 18 red algal genomes we analyzed, the same repeat annotation approach as outlined in the gene annotation section (**see Fig. 3A**). In the case of the male genome assembly of *Bostrychia*, the initial *de novo* repeat library was further manually curated to improve TE classification by both the modification of TE annotations and the structural refinement of selected consensus sequences. Each consensus sequence in the repeat library was compared to the eukaryotic repetitive sequences in the RepBase database v29_02^164^, using BLASTx^161,162^. The TE annotation of each consensus sequence was manually reviewed against the assigned RepBase annotation and adjusted in cases of strong e-value support. Selected TE families were also manually curated following^165^. In the initial TE library, the most abundant families were classified as *Mutator*-like DNA transposons (*MULEs*). To verify and refine this classification, the homology of several of these families was assessed by sequence alignment, which resulted in five distinct curated families (**Supplemental Fig. 13A**). Based on homology to transposase proteins, terminal sequence motifs and target site duplication lengths, these five families were re-classified as *EnSpm* superfamily TEs of the *Plavaka* lineage. The original misclassification stemmed from homology between the *Plavaka* accessory genes found both in *Bostrychia Plavaka* elements and *Chondrus crispus Mutator* elements (**Fig. 3A**). In addition, the consensus sequence of one abundant family belonging to LTR *Gypsy* elements was manually curated. To visualize the final TE complement in *Bostrychia*, a landscape of TE divergence was generated based on the percent divergence from the consensus sequence of their assigned family, which was derived from the RepeatMasker output (**Fig. 3B; Supplemental Fig. 13B**). TE and gene density was visualized using a Circos plot generated with the R package circlize (**Supplemental Fig. 8**)^166^.

To further analyze the structure of the *Plavaka* elements in *Bostrychia*, open reading frames of the accessory genes were identified with ORF finder using the manually curated consensus sequences^167^ (**Supplemental Fig. 13A**). InterPro was applied to functionally classify the accessory genes based on the presence of known protein domains^147^. For *Plavaka* accessory gene 2 (PAG2), a DNA methylase domain (PF01555) and a SET domain (PF00856) were identified. In contrast, no domains were detected for PAG1 or PAG3. To further investigate potential protein homology of these two accessory genes, HHpred searches were conducted using selected algal proteins identified through tBLASTn (using the *evalue* 0.001 option) with the respective PAG of *Bostrychia* as the query^168,169^. While no domain was identified for PAG1, analysis of PAG3 revealed a DNA-binding domain (best hit: 7BNR_Band, e-value 0.29) and a putative 2OG-dependent oxygenase domain (best hit: 2OPW_A, e-value 0.016).

Since it was infeasible to assign *Plavaka* elements through manual curation of the TE libraries across all investigated red algal genomes, an overlap-based approach was used as an alternative. Homologs of the *Bostrychia* PAG genes were first identified in each red algal TE library with tBLASTn (using the *evalue* 0.001 option). The PAG2 gene hits in each species were then used to determine whether the *Plavaka* transposase and the other two PAG genes lied within a 15 kb flanking region (total region 30 kb) on the same consensus sequence, to ensure that only complete *Plavaka* elements were recovered. The results of this analysis were validated by manual curation of selected families of *Plavaka* and other PAG-associated elements in *Porphyra umbilicalis, Pyropia yezoensis, Gracilaria chilensis* and *Kappaphycus alvarezii* (**Supplemental Table 6**)

### Gene and TE expression analysis

The RNA-seq samples of female gametophyte, male gametophyte and tetrasporophyte were processed and gene expression levels analyzed using the nf-core RNA-seq pipeline v3.12.0-g3bec233 (nextflow v23.10.0.5889) with Salmon as the alignment tool^170,171^. Differential gene expression analysis was performed using DESeq2 v1.40.2^172^. Statistical tests for statistical comparisons of gene expression were performed using the rstatix package in R^173^. The expression of TEs in *Bostrychia* was quantified using TEspeX v2.0.1 using the female, male and tetrasporophyte RNA-seq samples^174^. Tandem repeats together with annotated rRNA sequences were used as non-coding transcripts. Expression levels of TE families were quantified as TPM values by normalizing the raw counts from the TEspeX output to their respective consensus sequence lengths.

### Genome size and ploidy estimation

DNA content and ploidy levels of *Bostrychia moritziana* (male and female gametophytes) (JW3235)*, Bostrychia montagnei* (JW2927)*, Bostrychia flagellifera* (JW3553)*, Bostrychia scorpioides* (JW4873)*, Caloglossa monosticha* (male gametophyte) (JW4013) and *Polysiphonia senticulosa* (KU-0950) were estimated with flow cytometry using *Zea mays L. ‘CE-777’* as the DNA reference standard (**Supplemental Table 2**)^175^. To remove potential surface contaminants, including microbial communities and residual tissue, algal tissue was cleaned in NSW with the addition of sand and vortexed using a Precellys Evolution beads homogenizer (Bertin technologies) three times. 20 mg of plant tissue and 20 mg of the cleaned dried algal tissue were placed in the center of a petri dish on ice. 500 µl of nuclei isolation buffer (NIB) (15 mM Tris-HCl pH 8, 2 mM EDTA pH 8, 80 mM KCl, 20 mM NaCl, 0.04% 2-Mercaptoethanol, 0.5 mM Spermine, 0.1% BSA, 0.1% Triton X-100) was added and the tissues chopped using a razor blade for 2 minutes. An additional 500 µl of NIB buffer was then added to the homogenate and the nuclei solution was filtered through a 40 µm filter. The nuclei were pelleted by centrifugation at 500 × g for 5 minutes at 4°C. The pellet was gently resuspended in 500 µl of NIB buffer and 2 ml of staining solution (Cystain PI Absolute P, Sysmex, Product 05-5022) was added. Before loading onto the flow cytometer, 1 ml of the stained nuclei solution was diluted with 1 ml of staining solution. The samples were measured using a BD FACSMelody Cell Sorter (BD Biosciences) and the measurements further analyzed using FlowJo v10.10.0^176^. To search for signs of a potential whole genome duplication in *Bostrychia,* wgd2 using the *wgd dmd, ksd* and *viz* parameter was used^177,178^. Nucleotide-based self-synteny dot plots for the male and female assembly were generated using D-Genies^179^ (**Supplemental Fig. 10C&D**).

### Phylogenetic trees and ancestral state reconstructions

Four different phylogenetic trees were constructed, each tailored to address different questions and applications. To perform the ancestral state reconstruction of genome size estimates from Kapraun & Freshwater (2012), a concatenated gene tree was generated using conserved plastid, mitochondrial and nuclear marker genes (atpA, atpB, cox1, psaA, psbA, rbcL and EF2) from red algae and selected outgroup species (**Supplemental Table 1**). Gene sequences were retrieved from the NCBI Protein database by searching with the keywords “Rhodophyta” and the respective gene name^180,181^. The dataset was then filtered to include only species for which at least three of the seven marker genes were present. Individual gene alignments were generated using MAFFT v7.520 using the *-auto* option and subsequently concatenated with a custom python script^182^. ModelTest-NG v0.1.7 was used to determine the optimal model of evolution per gene partition^183^. A maximum likelihood phylogeny was inferred by first running RAxML (raxmlHPC-PTHREADS-AVX) using the PROTGAMMALG amino acid substitution model. Rapid bootstrap support was enabled with the –k option, the –# and –x options using 100 replicates. Bootstrap values were assigned in a second run using the –z option^184^. The species names in the phylogeny were matched to those with available genome size estimates from Kapraun & Freshwater (2012). In the absence of an exact match, species names were adjusted to correspond to another species within the same genus in the phylogeny (**Supplemental Table 1**). To find the appropriate model of trait evolution, we iterated over 100 random bootstrap replicates using the function *fitDiscrete* implemented in the Geiger package v2.0.11^185^. The corrected Akaike information criterion was used to define the Brownian motion model as the most appropriate model. To reconstruct the ancestral states, the function *anc.ML* was used and the states were mapped using the functions *contMap* and *setMap* implemented in the phytools package v2.3.0^186^.

To perform the analysis of expanded and contracted gene families, a species tree based on 2,372 conserved OGs across red algae and the green lineage was constructed. Protein sequences of 39 species (**Supplemental Table 3 DS2**) were grouped into 45,785 orthogroups using OrthoFinder v2.5.5 with the *-m* and *-x* options^144^. OGFilter was applied to retain OGs containing sequences from at least 80% of the species^145^. ParGenes v1.2.0 was used for best-fit model selection and gene tree inference for each of the 2,372 filtered OGs, using the *-m* and the *--use-astral* options^187^. The final species tree was generated using Astral-Pro v1.16.1.3, rooted with the cryptophyte *Guillardia* as the outgroup and ultrametricized using the *make_ultrametric.py* script from OrthoFinder^145,189,190^. Before performing the final gene family analysis, the tree was pruned to include only red algal species and the outgroup species. In addition, the same phylogeny was used to perform the ancestral state reconstruction of *Plavaka* elements, treating their presence as a discrete trait. The appropriate model of trait evolution was assessed as described above, with the Early-burst model selected for analysis. Ancestral states were reconstructed and plotted onto the phylogeny using the *rayDISC* and *plotRECON* functions from the corHMM package v2.8^191^.

For the analysis of *EnSpm* DNA transposons, a phylogenetic tree was constructed using 335 eukaryotic *EnSpm* transposase proteins, with *Chapaev* transposases serving as the outgroup. The dataset included sequences from the RepBase database v29_02, manually curated red algal *Plavaka* transposases and the original fungal *Plavaka* transposases from^45^. Proteins with at least 60% sequence similarity over at least 60% of their length were reduced to a single representative using CD-HIT v4.8.1 (*-d 0 –aS 0.6 –c 0.6 –G 0 –g 1 –n 3*)^192^. Sequence alignment was performed using MAFFT v7.520 (“L-INS-i”), gappy columns were removed using trimAl (*-gappyout*)^183^, and finally the alignment was manually trimmed at each end to capture the DDE catalytic domain^193^. The maximum likelihood phylogeny was inferred using IQ-TREE v2.2.2.7, using ModelFinder for best-fit model selection (*-MFP*), and support values were assessed with ultrafast bootstrapping and SH-aLRT using 1000 replicates (*-alrt 1000, –bb 1000*)^194^.

A total of 159 protein sequences were collected for the maximum likelihood phylogeny of TALE-HD TFs, based on annotations from TAPscan for species of the green lineage, Cryptophyta, brown algae and red algae. The alignment was generated with MAFFT v7.520 with the *-auto* option, and alignment trimming was conducted using trimAl v1.4.rev15 to remove gappy columns using the –gt 0.1 option^183,195^.

### Comparative genomic analyses

Gene ages in *Bostrychia* were computed using GenEra v1.4.0 (**Supplemental Table 7**)^196^. OrthoFinder v2.5.5 was used with the *-m* and *-x* options to group *Bostrychia* protein sequences with those of 19 other red algal species into 20,618 orthogroups (OGs) (**Supplemental Table 3 DS3**)^145^. Ceramiales– and Rhodomelaceae-specific OGs were identified by determining overlaps using the R package UpSetR^197^. Expansions and contractions in OG sizes were estimated using CAFE5 v5.1 under a birth and death model using the OG counts of the 20 red algal species defined by OrthoFinder. The species tree was constructed from conserved OGs as described in the phylogenetic tree methods section^198^. In the first run, we estimated the initial birth-death parameter (λ). In the second run, we calculated an error model to account for assembly and annotation errors using the *-e* option. In the third run, the error model was applied to refine the estimation of the final λ parameter. Finally, in the fourth run, both the error model and the final λ parameter were applied. To further analyze the expanded and specific OGs in the Ceramiales and Rhodomelaceae, we performed a Pfam domain enrichment analysis by comparing domain frequencies in the subset of expanded genes compared to the background (all species) using Fisher’s Exact Test in R. p-values were adjusted for multiple testing using the Benjamini-Hochberg method to identify significantly enriched domains (adjusted p-value < 0.05).

### Annotation of carbohydrate-active enzymes (CAZymes)

Predicted GTs and GHs were determined from the *Bostrychia* proteome using dbCAN3 v4.1.4 (protein mode, default settings)^199^. *Bostrychia* proteins were filtered by requiring identification by ≥2 CAZyme search tools within dbCAN3 (HMM, dbCAN_sub, DIAMOND) or a single tool hit with additional familial matching against domains detected via InterProScan v.5.67-99.0^200^. Identified families were assigned by the highest scoring match using BLASTp (default settings)^162,163^ against datasets comprising characterised CAZyme protein sequences from each GH and GT family^201^ [http://www.cazy.org/] (accessed October 2024). Orthogroups were produced using OrthoFinder v2.5.5 (default settings)^145^ and cross-analysed with dbCAN3 outputs to determine orthology within GH and GT families. Protein sequence alignment performed using MUSCLE (default settings) and phylogeny reconstruction with Maximum likelihood method (bootstrap method, 500 bootstrap replicates) using MEGA11^202^. To identify CAZymes across other red algal species, a CAZyme protein was considered present if it had at least one representative sequence within the same orthogroup as a reference CAZyme of *Bostrychia*.

### Sex chromosome detection

To determine the structure of the sex chromosomes in *Bostrychia*, we employed a k-mer-based approach to reciprocally map reads from one sex onto the assembly of the opposing sex using the YGS method^83,88,203^. The male and female HiFi reads were split into 150 bp reads using SeqKit v2.5.0^204^. Jellyfish v2.3.1 count and dump were used to split these reads into canonical k-mers of size 15, using the parameters *quality-start* 33, *min-quality* 20, and *lower-count* 5^205^. Both the male and female assemblies were divided into 50 kb-sized chunks using SeqKit. Eventually, the YGS method as described in Carvalho & Clark (2013) was used to map k-mers of one sex onto the fragmented assembly of the other sex. Regions with a high density of unmatched single-copy k-mers, which are indicative of sex-linked regions, were only observed on one chromosome in both the male and female gametophyte (**Supplemental Fig. 18A&B**). These putative sex chromosomes were among the smallest chromosomes in *Bostrychia*, measuring out at 19.98 Mb and 20.87 Mb in the female and male, respectively (**Fig. 5A**). The first portion of these chromosomes had high mapping coverage and is indicative of a PAR, spanning 7.61 Mb in the female and 7.74 Mb in the male. Downstream of the PAR is a region with over 80% unmapped reads, highlighting an extended sex-specific region representing the female (12.37 Mb) and male SDR (13.13 Mb) respectively (**Fig. 5A**).

Synteny between the sex chromosomes and the identification of gametologs was assessed using reciprocal best hit BLAST searches (BLASTn using *-max_target_seqs* 1 and *-max_hsps* 1). Synonymous substitution rates were computed as a proxy for the relative age of gametologs. Protein alignments of the gametolog pairs were first created using MAFFT v7.520 with the *--auto* option, and codon-based alignments were then generated by mapping the protein alignments to the CDS using pal2nal v14 with the *-nogap* option^206^. Finally, CODEML out of the PAML package v4.10.7 was used to calculate dS values for each gametolog pair^207^. To determine the presence of gametologs in other red algal species, a gametolog was considered present in a species if it was assigned to the same orthogroup as the respective gametolog in *Bostrychia.* The genes on the sex chromosomes were manually grouped into 11 functional classes based on their assigned functional annotations (**Supplemental Table 9**). Annotation of TALE-HD transcription factors across six phyla was performed using TAPscan v4^151^. To further classify red algal TALE-HD TFs into the KNOX-Red1, KNOX-Red2, and PBX classes, *hmmbuild* from the HMMER v3.4 software package was used to construct profile hidden Markov models (HMMs) for the KN-A, KN-B, and PBL domains based on alignments provided in Joo et al., 2018^208^. These HMMs were then used with *hmmsearch* to identify the corresponding domains across the red algal TFs. Additionally, the KNOX-Red1 and KNOX-Red2 sequences from Joo et al., 2018 were aligned separately using MAFFT v7.520 with the *--auto* option, with the conserved homeodomain removed. Profile HMMs were then constructed and used for searches following the same approach as for the other domains.

To further confirm sex-specific expression and conservation of UV genes, we used previous data from a geographically-distinct strain of *Bostrychia* (JW2746)^26^. We used primer sequences from Shim et al. (2021) that were designed to amplify sex-specific genes by genomic PCR as query sequences for BLASTn searches (*-word_size 7* option) against coding sequences on the male and female sex chromosomes of *Bostrychia* (**Supplemental Table 9**). In addition, we analyzed gene expression levels using the provided RNA-seq samples from Shim et al. (2021) of the female gametophyte, male gametophyte and tetrasporophyte as described above (**Supplemental Table 9**).

## Data and material availability

The *Bostrychia moritziana* v1.0 genome assembly has been uploaded to the National Center for Biotechnology Information (NCBI) and is available under BioProject PRJNA1219826. The raw ONT and PacBio long-reads and Illumina Hi-C and RNA-seq short-reads have also been uploaded to BioProject PRJNA1219826. A repository containing phylogenetic trees, manually curated TE libraries, the female *Bostrychia* assembly and annotation, and filtered genome assemblies and annotations for selected red algal species can be accessed at https://doi.org/10.17617/3.RTULYR. All scripts and red algal cultures used in this study are available upon request from the corresponding author.

## Author contributions

Conceptualization, MB; Methodology, RP and MB; Investigation, RP, OM, MO, RC and MB; Visualization, RP and MB; Resources, JAW and SMC; Data Curation, RP; Writing – Original Draft, RP, OM, MO and MB; Writing – Reviewing & Editing, RP, JW, OM, MO, RC, SMC and MB; Supervision, MB; Project Administration, MB.

## Acknowledgments

We thank Alan Critchley for originally collecting *Bostrychia* specimens used in this study, Masakazu Hoshino for advice on algal culture, Elena Avdievich for support during Nanopore sequencing, Pengfei Liu for support during Hi-C analysis, Julia Golebiowska for assistance during RNA-seq library preparation and Agnieszka Lipinska for useful discussions and guidance for sex chromosome detection. RP is a member of the International Max Planck Research School ‘From Molecules to Organisms’. RP, MB, and SMC were supported by the Max-Planck-Gesellschaft. MO was supported by a Novo Nordisk Foundation Project Grant (NNF21OC0071501).

## Supplementary Figures

Supplemental Figure 1. Genome size estimates across red algae.

Supplemental Figure 2. Rooted phylogenetic tree of red algae based on marker genes.

Supplemental Figure 3. Flow cytometry results of species belonging to the Ceramiales.

Supplemental Figure 4. Karyoplot of *Bostrychia* highlighting telomeres, rRNA, and gaps.

Supplemental Figure 5. Mitochondrial and chloroplast genomes of *Bostrychia*.

Supplemental Figure 6. Gene expression patterns across *Bostrychia* life stages.

Supplemental Figure 7. Comparative analysis of gene features in *Bostrychia* and other red algae.

Supplemental Figure 8. Circos plot of gene and repeat distribution across the *Bostrychia* genome.

Supplemental Figure 9. Flow cytometry results of *Bostrychia*.

Supplemental Figure 10. Assessment of a possible whole genome duplication event.

Supplemental Figure 11. Alignment of *Plavaka* transposase sequences.

Supplemental Figure 12. Rooted phylogenetic tree of eukaryotic *EnSpm* transposase proteins.

Supplemental Figure 13. Structure and coverage of *Plavaka* elements across red algae and TE expression in *Bostrychia*.

Supplemental Figure 14. Gene age distribution across the *Bostrychia* genome.

Supplemental Figure 15. Annotated Pfam domains in *Bostrychia* genes.

Supplemental Figure 16. Genome browser view of selected gene clusters in *Bostrychia*.

Supplemental Figure 17. Unrooted phylogenetic trees of mannosylglycerate hydrolases and β-1,4-mannanases from red algae.

Supplemental Figure 18. Identification and analysis of sex chromosomes in *Bostrychia.* Supplemental Figure 19. Alignment of TALE-HD transcription factors across the Archaeplastida. Supplemental Figure 20. Blob plots of the *Chondria, Digenea* and *Griffithsia* draft assemblies.

## Supplemental Tables

Supplemental Table 1. Genome size estimates, phylogenetic tree data, and adjustments for ancestral state reconstruction.

Supplemental Table 2. Results of flow cytometry DNA content estimations in red algae.

Supplemental Table 3. Species datasets, sequencing and assembly information.

Supplemental Table 4. Genome expression data.

Supplemental Table 5. Structural and functional annotation data.

Supplemental Table 6. Manually curated transposable elements and repeats that carry at least one *Plavaka* associated gene.

Supplemental Table 7. Results of gene age inference and Pfam domain enrichment analysis.

Supplemental Table 8. Annotated carbohydrate-active enzymes.

Supplemental Table 9. Annotation and expression analysis of sex chromosomes.

## Notes

### Competing Interest Statement

The authors have declared no competing interest.

