## Supplemental Figures for "The expanded *Bostrychia moritziana* genome unveils evolution in the most diverse and complex order of red algae"

### Supplementary Figures

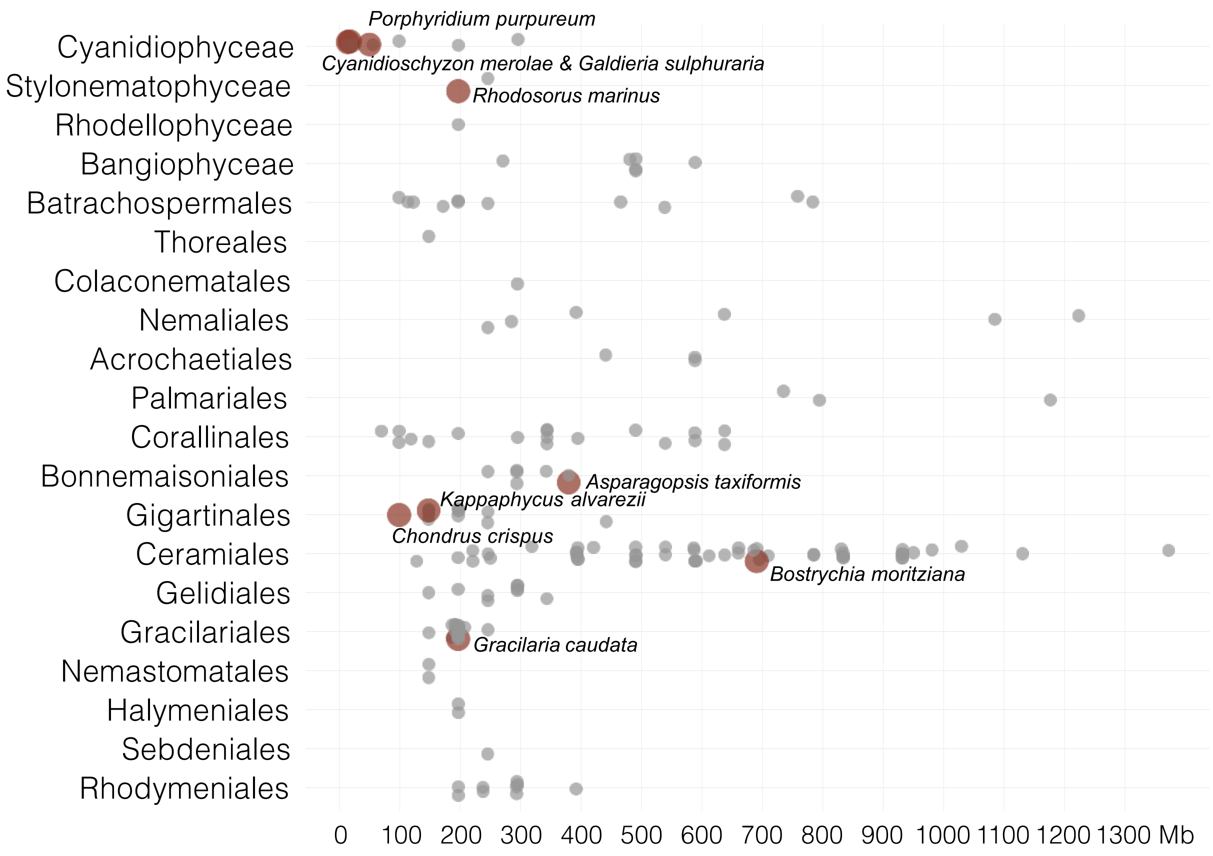


**Supplemental Figure 1. Genome size estimates across red algae.**
A dot plot showing genome size estimates (converted to Mb) from Kapraun & Freshwater, 2012 on the x-axis and taxonomic classification on the y-axis. Species are grouped by class, with orders displayed for the Florideophyceae. Each point represents the genome size estimate of each species, with those with a published genome assembly used in this study highlighted in red.

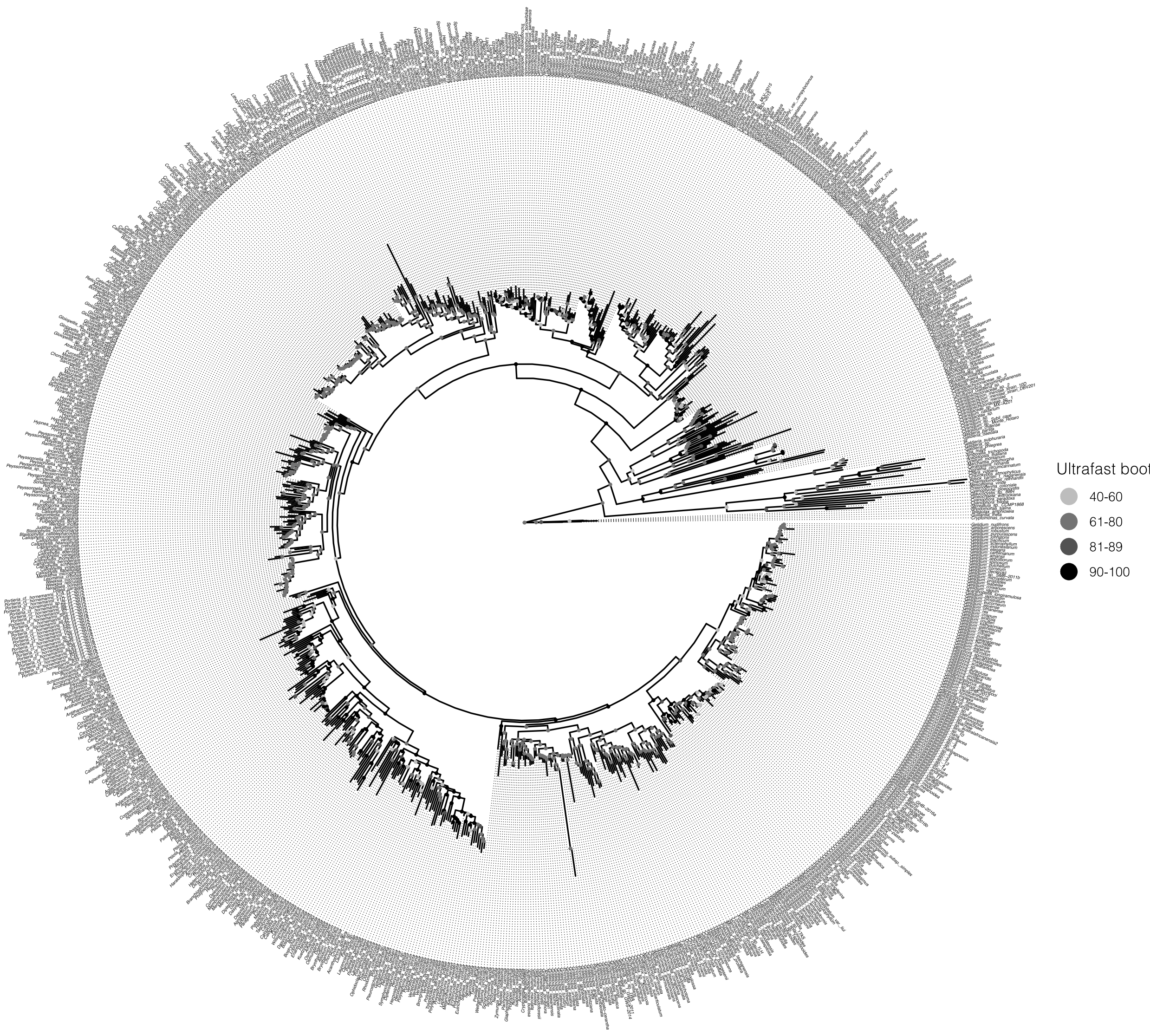


**Supplemental Figure 2. Rooted phylogenetic tree of red algae based on marker genes.**
A phylogenetic tree inferred from concatenated plastid, mitochondrial, and nuclear marker genes (atpA, atpB, cox1, psaA, psbA, rbcL, and EF2) of 939 red algal and selected outgroup species (species of the green lineage, Glaucophytes, and Cryptophytes). Cryptophyte sequences were used as the root. Ultrafast bootstrap values are indicated as color-coded dots at each node of the phylogeny.

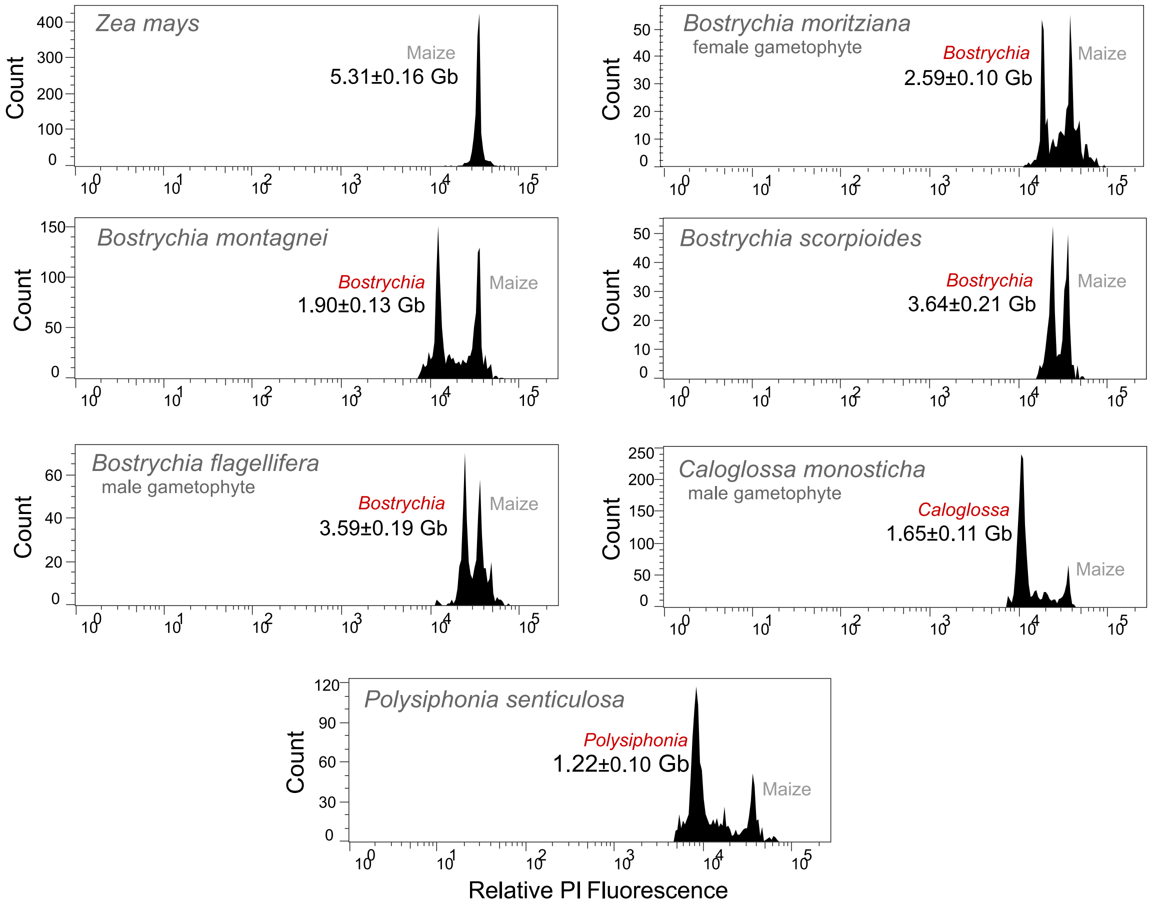


**Supplemental Figure 3. Flow cytometry results of species belonging to the Ceramiales.**

Flow cytometry analysis of propidium iodide-stained nuclei from *Bostrychia moritziana, Bostrychia montagnei, Bostrychia scorpioides, Bostrychia flagellifera, Caloglossa monosticha* and *Polysiphonia senticulosa* alongside nuclei isolated from leaf tissue of *Zea mays.* The estimated DNA content (in Gb) is indicated alongside the red algal peaks.

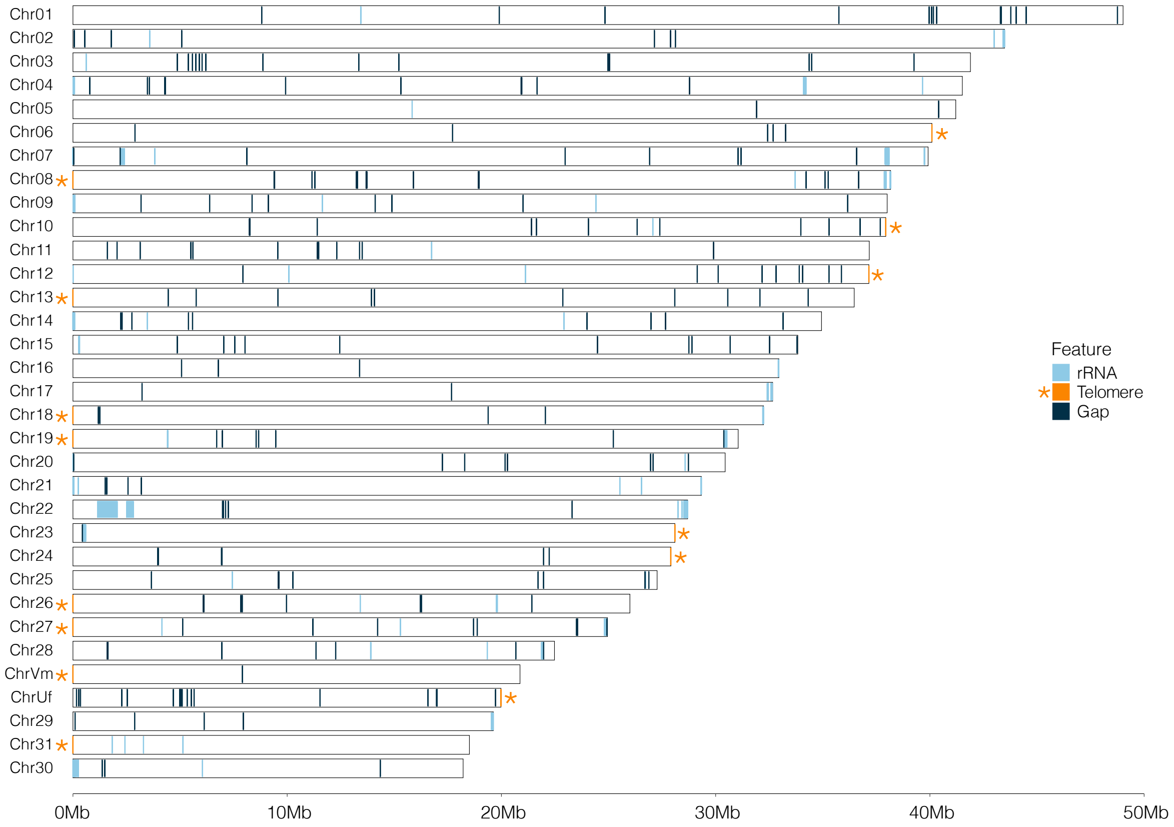


**Supplemental Figure 4. Karyoplot of *Bostrychia* highlighting telomeres, rRNA, and gaps.**
A karyoplot displaying all 31 autosomal chromosomes of *Bostrychia*, along with the female (U) and male (V) sex chromosomes. A scale bar indicating chromosome size (in Mb) is shown below the chromosomes. rRNA clusters are shown in light blue, gaps in dark blue, and telomeres in orange. Orange asterisks further highlight telomere locations.

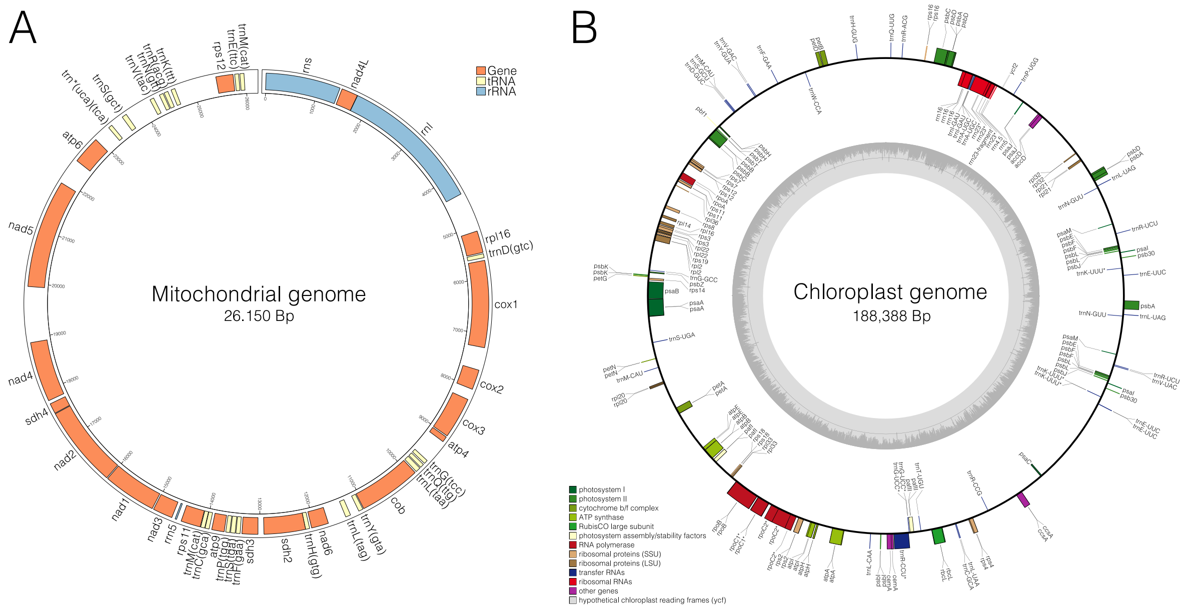


**Supplemental Figure 5. Mitochondrial and chloroplast genomes of *Bostrychia*.**
(**A**) Circular mitochondrial genome of *Bostrychia* with gene, tRNA and rRNA annotations. Genes are shown in orange, tRNAs in yellow and rRNA in blue. (**B**) Circular chloroplast genome with color-coded functional categories: photosynthesis genes in green, RNA polymerases in dark red, ribosomal proteins in brown, tRNAs in dark blue, rRNA in light red, other genes in purple, and hypothetical chloroplast reading frames in grey.

A

| Sample | Reads | %Aligned | %Duplicate reads |
|---|---|---|---|
| Female rep 1 | 20,440,515 | 88.0 | 65.35 |
| Female rep 2 | 20,950,427 | 83.3 | 65.55 |
| Female rep 3 | 18,184,343 | 85.5 | 64.10 |
| Male rep 1 | 17,263,056 | 84.0 | 62.10 |
| Male rep 2 | 16,420,553 | 76.2 | 67.55 |
| Male rep 3 | 25,399,539 | 81.2 | 66.70 |
| Tetrasporophyte rep 1 | 20,597,235 | 78.9 | 65.85 |
| Tetrasporophyte rep 2 | 22,179,758 | 66.6 | 71.70 |
| Tetrasporophyte rep 3 | 20,758,948 | 80.7 | 63.25 |

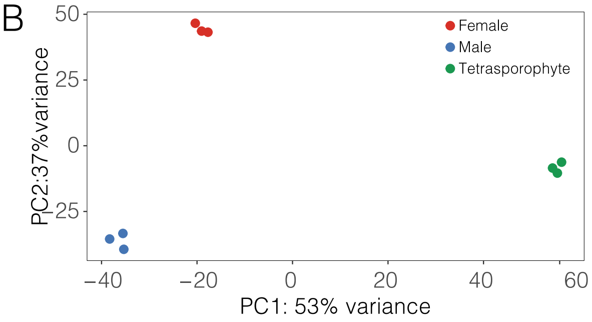


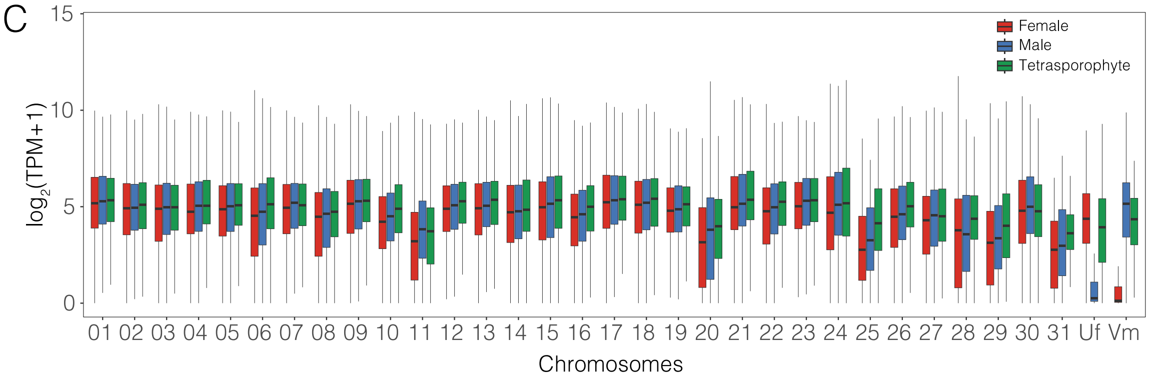


**Supplemental Figure 6. Gene expression patterns across *Bostrychia* life stages.**
(**A**) Table summarizing the number of reads, percentage of aligned reads and the percentage of duplicated reads for each dataset. (**B**) Principal component analysis (PCA) illustrating variance between replicates of transcriptomic datasets used in study. Each stage was profiled with three biological replicates. (**C**) Boxplot showing the distribution of log2 transformed TPM+1 values in the female gametophyte (red), male gametophyte (blue), and the tetrasporophyte (green) for genes grouped by chromosome.

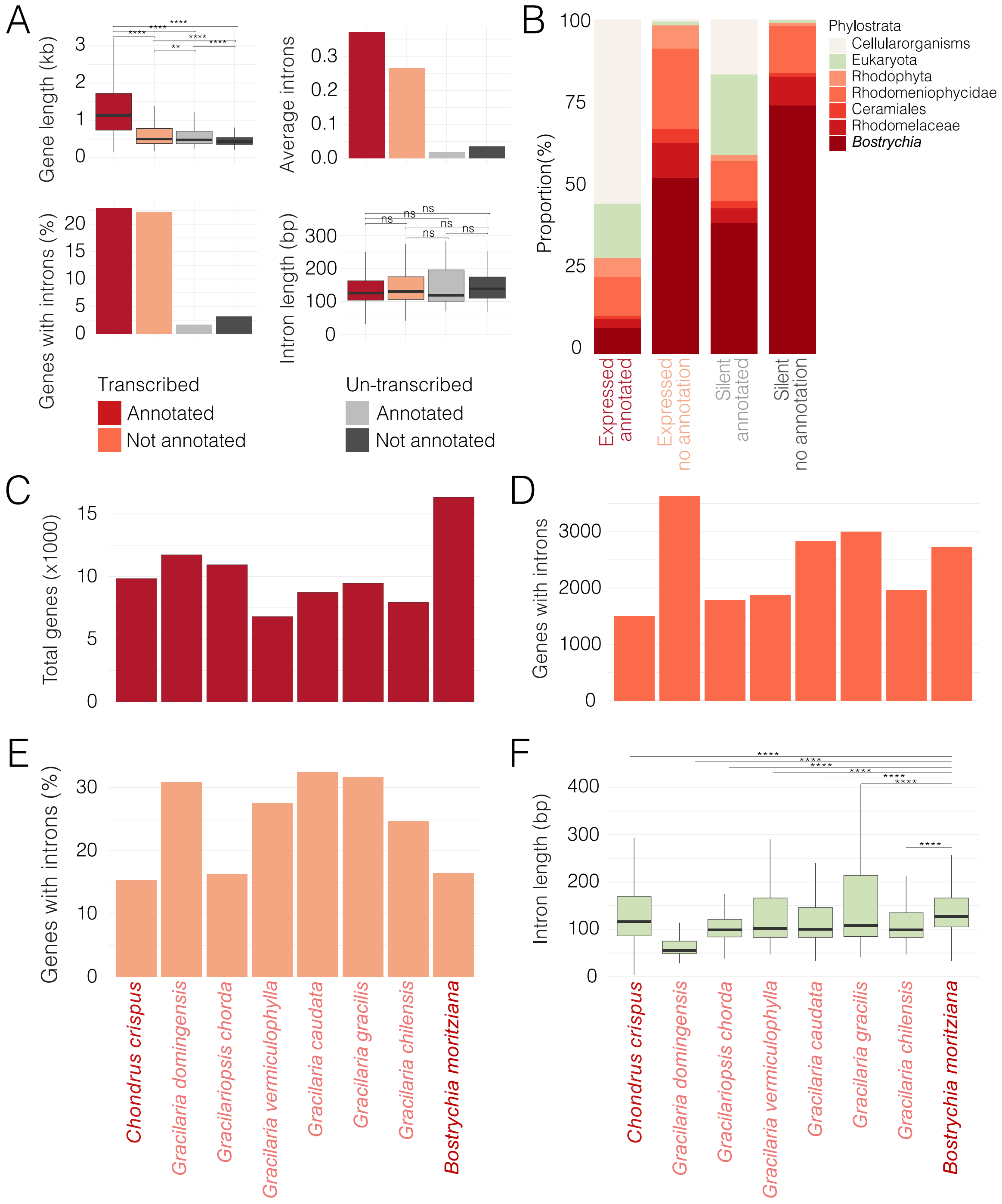


**Supplemental Figure 7. Comparative analysis of gene features in *Bostrychia* and other red algae.** (**A**) Gene length (in bp) of transcribed, untranscribed, annotated and unannotated genes in *Bostrychia*, alongside the average number of introns per gene, the percentage of introns, and the average intron length (in bp) for each group. (**B**) Phylostratigraphic distribution of gene age across (un-)transcribed and (un-)annotated genes. (**C-F**) Total number of genes (**C**), number of genes containing introns (**D**), percentage of genes with introns (**E**), and average intron length (in bp) (**F**) for a selection of Florideophyceae species. Wilcoxon test results for statistical comparisons are shown above the boxplots. * $p$-value < 0.05, ** $p$-value < 0.01, *** $p$-value < 0.001, ns = no significance.

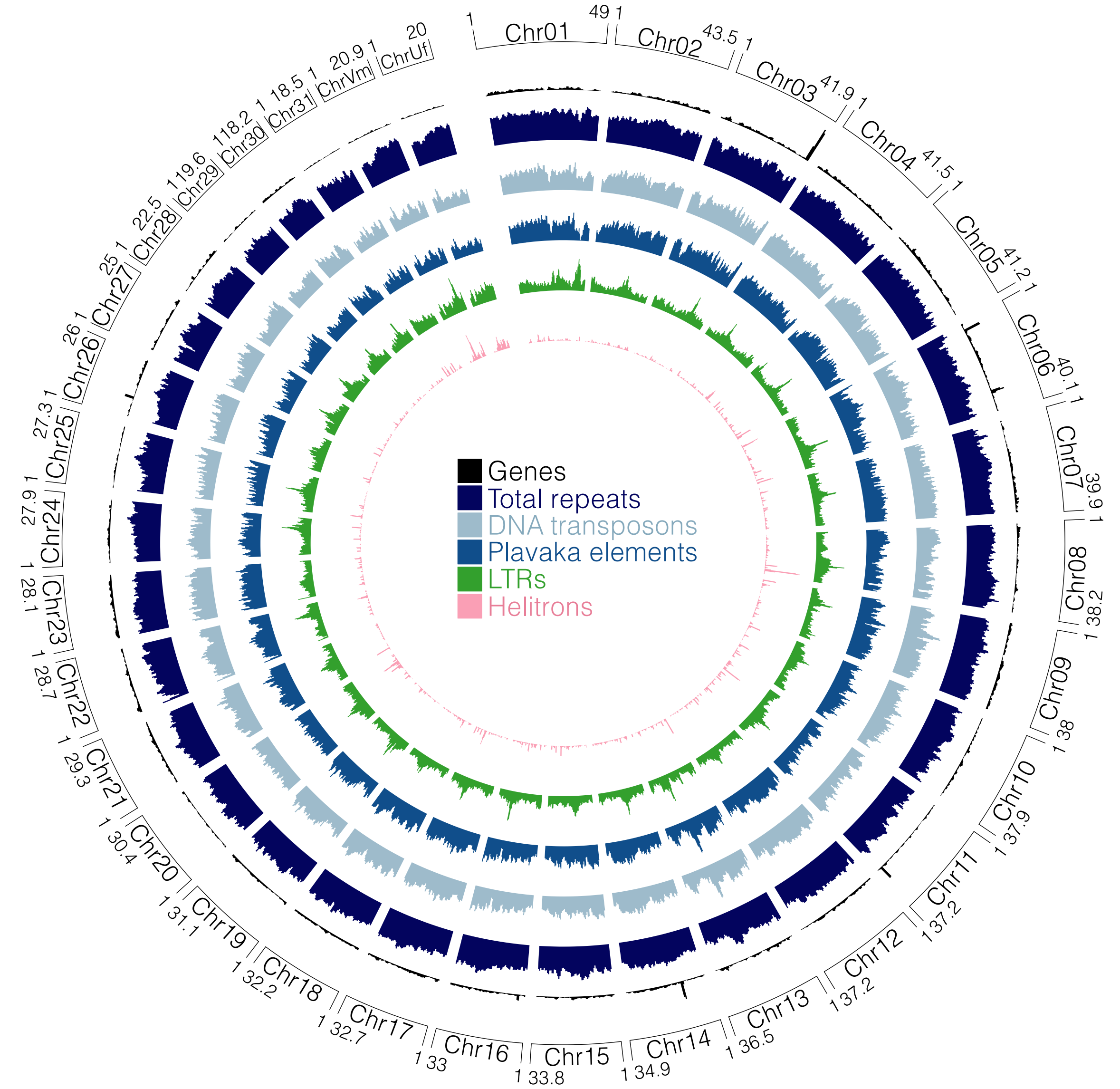


**Supplemental Figure 8. Circos plot of gene and repeat distribution across the *Bostrychia* genome.** Circos plot showing gene density (black), total repeat density (dark blue), density of DNA transposons (light blue), density of *Plavaka* elements (medium blue), LTR density (green) and density of Helitrons (pink) across all chromosomes in *Bostrychia.* Chromosome lengths are indicated next to the chromosome names.

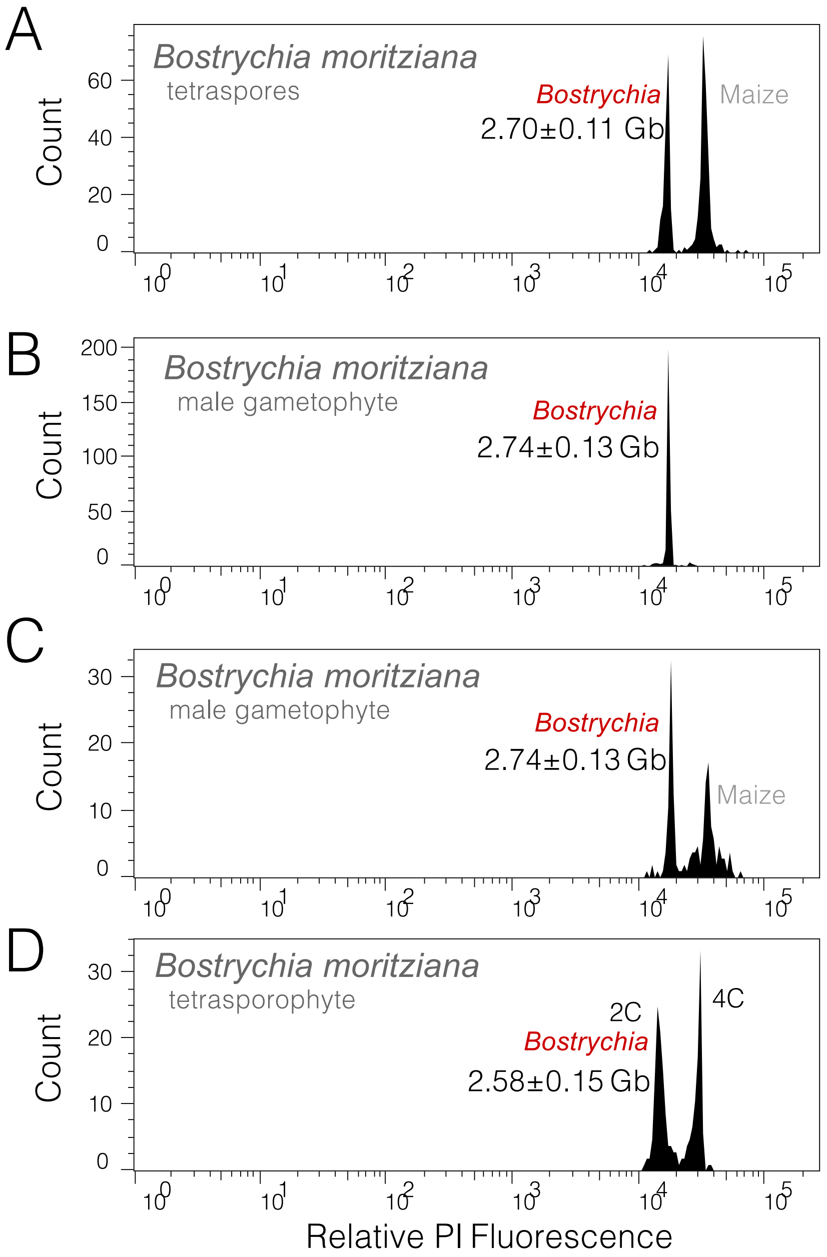


**Supplemental Figure 9. Flow cytometry results of *Bostrychia*.**
Flow cytometry analysis of propidium iodide-stained nuclei of tetraspores of *Bostrychia* alongside nuclei isolated from leaf tissue of *Zea mays* (**A**), the male *Bostrychia* gametophyte (**B**), the male *Bostrychia* gametophyte alongside *Zea mays* (**C**), and the tetrasporophyte of *Bostrychia* alongside *Zea mays* (**D**). The estimated DNA content (in Gb) is indicated alongside the red algal peaks.

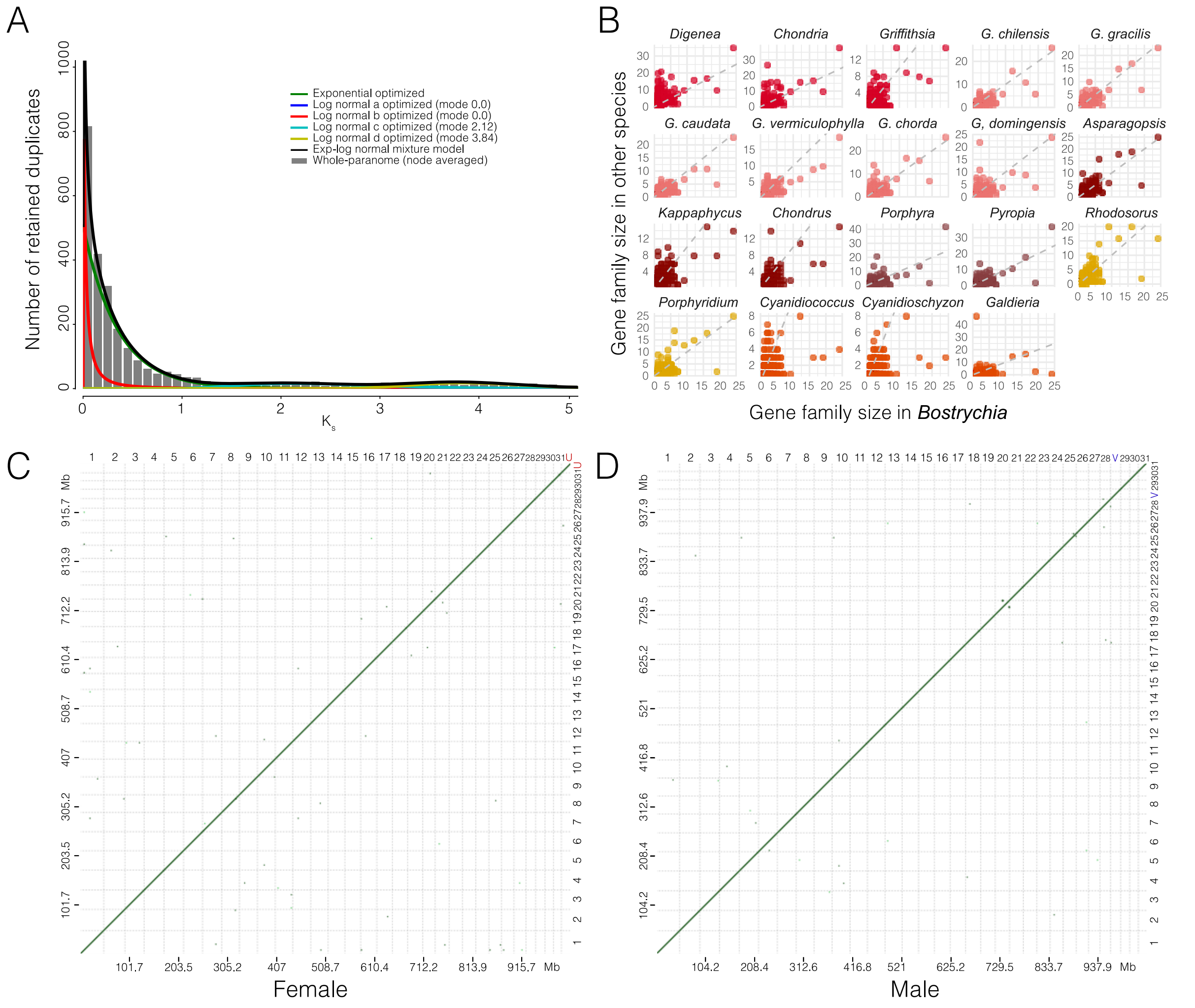


**Supplemental Figure 10. Assessment of a possible whole genome duplication event.**
(**A**) Distribution of synonymous substitution rates (Ks) for retained duplicate genes in *Bostrychia*, with multiple model fits overlaid. The absence of a distinct peak in the Ks distribution suggests no evidence of a recent whole-genome duplication event. (**B**) Dot plots comparing gene family sizes in orthogroups common to all investigated red algae between *Bostrychia* and other red algae. The color-code refers to the order of the respective red algal species. (**C-D**) Self-synteny dot plot of the *Bostrychia* female assembly (**C**) and male assembly (**D**).

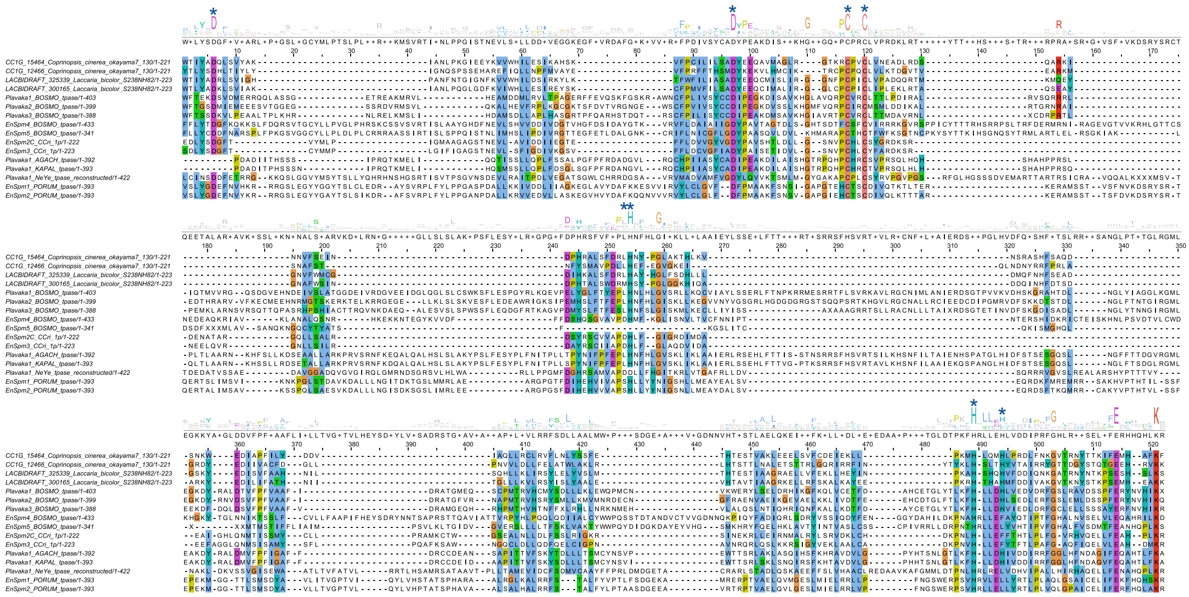

**Supplemental Figure 11. Alignment of *Plavaka* transposase sequences.**

Alignment of *Plavaka* transposase sequences from selected basidiomycete fungi, manually curated transposase sequences from *Bostrychia*, and other selected red algae, and sequences of *Chondrus crispus.* The color-code refers to conserved positions using the Clustal color scheme. The sequence logo represents conservation at each position. Dark blue asterisks highlight shared conserved residues of DDE type transposases according to Yuan & Wessler, 2011.

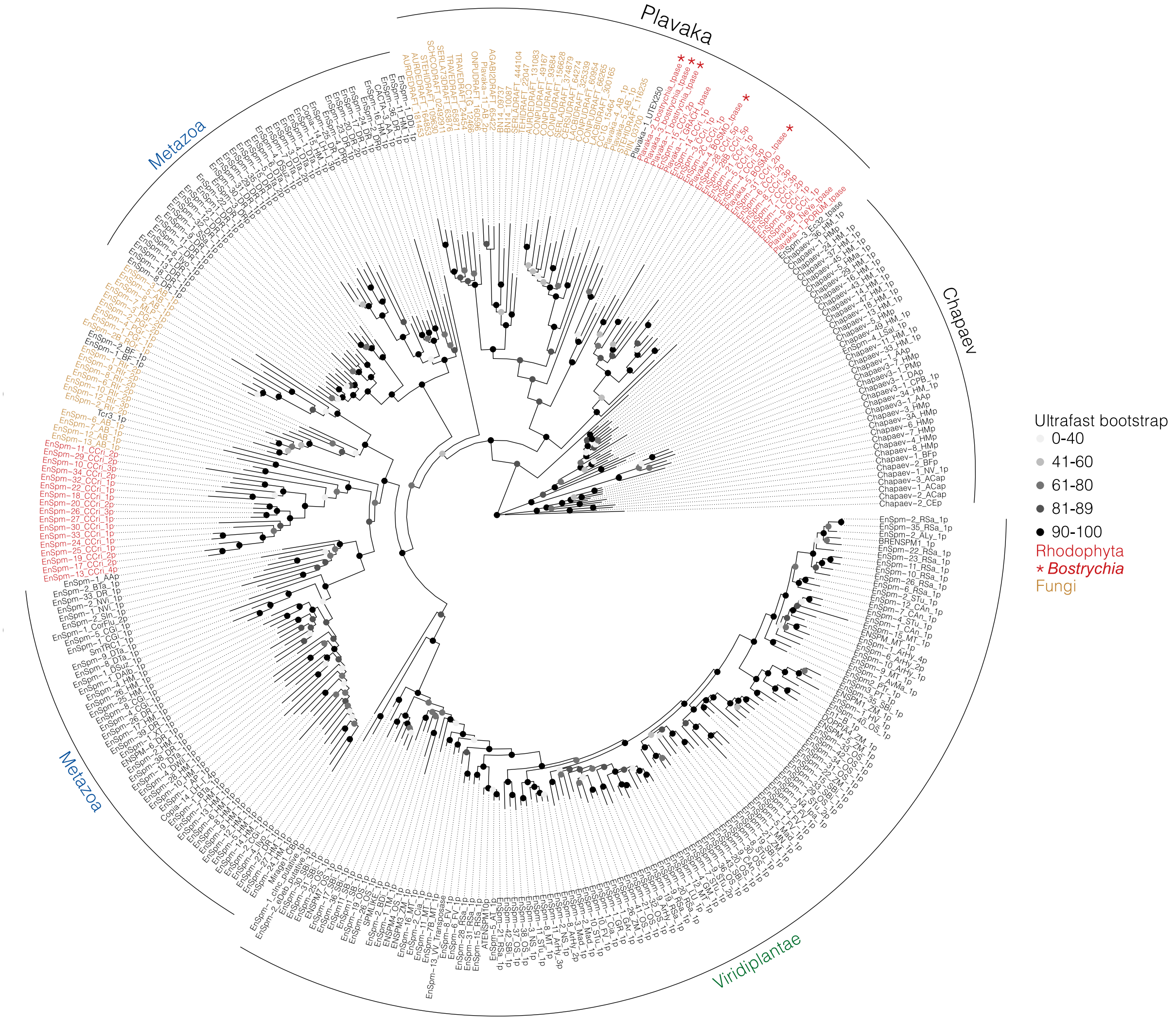


**Supplemental Figure 12. Rooted phylogenetic tree of eukaryotic *EnSpm* transposase proteins.** Phylogenetic tree inferred from 335 eukaryotic *EnSpm* transposase proteins of sequences from the RepBase database, manually curated red algal *Plavaka* transposases (in red, highlighted with asterisks) and the original fungal *Plavaka* transposases (in yellow). *Chapaev* transposase sequences were used as the outgroup. Ultrafast bootstrap values are indicated as color-coded dots at each node of the phylogeny.

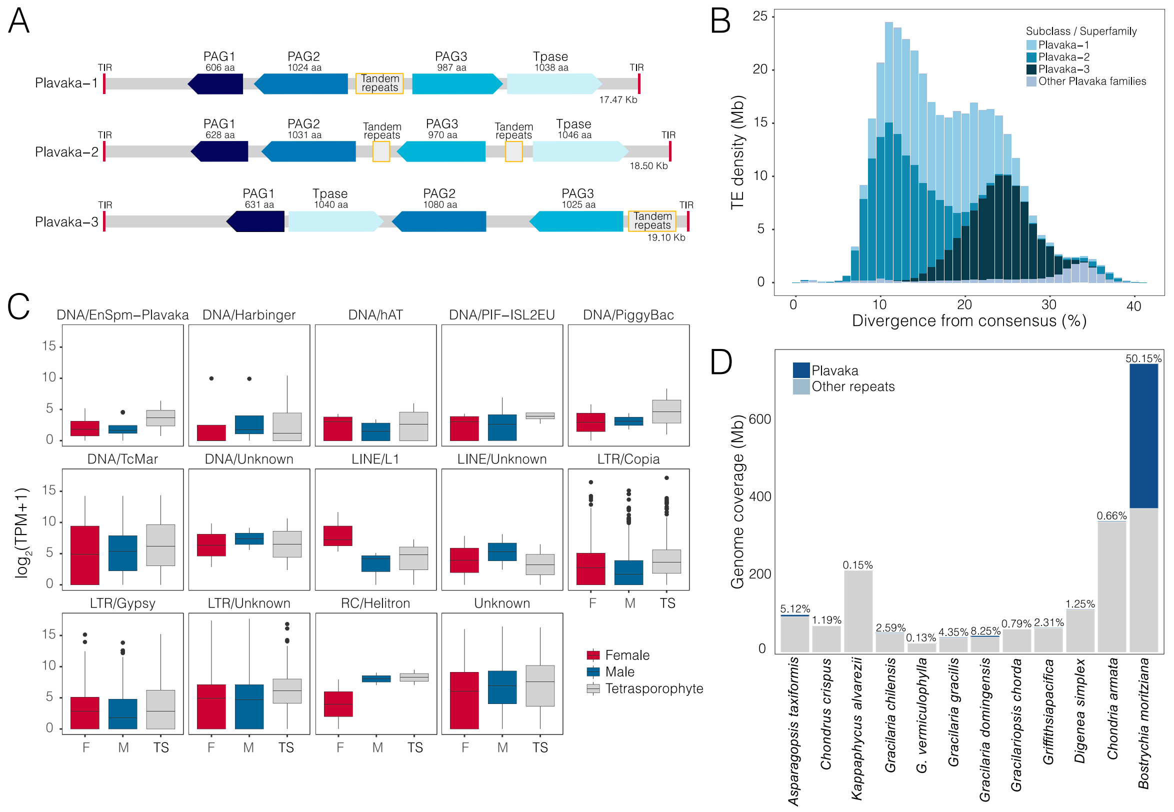


**Supplemental Figure 13. Structure and coverage of *Plavaka* elements across red algae and TE expression in *Bostrychia*.**

(**A**) Schematic representation of the *Plavaka* element structure in *Bostrychia* highlighting the relative position of the three *Plavaka*-associated genes (PAG1-3), the tandem repeats and the transposase gene of the *Plavaka-1*, *Plavaka-2* and *Plavaka-3* families. (**B**) The sequence divergence landscape of *Plavaka* elements in *Bostrychia*. TE density (in Mb) of each *Plavaka* element grouped by family is plotted against the divergence from the consensus sequence of its assigned family. (**C**) Boxplots showing the distribution of log2-transformed TPM+1 values from TEs in the female gametophyte (red), male gametophyte (blue), and the tetrasporophyte (grey) grouped by TE class. (**D**) Genomic coverage of repeats and *Plavaka* elements across different red algal species. The x-axis represents the species, while the y-axis indicates genomic coverage (in Mb). Each bar is subdivided to show the total repeat content in grey, with *Plavaka* elements highlighted in dark blue. The percentage above each bar represents the proportion of *Plavaka* elements relative to the total repeat content in each species. Due to the lack of manual curation of *Plavaka* elements in red algae other than *Bostrychia*, the percentage of *Plavaka* elements in these species may be underestimated.

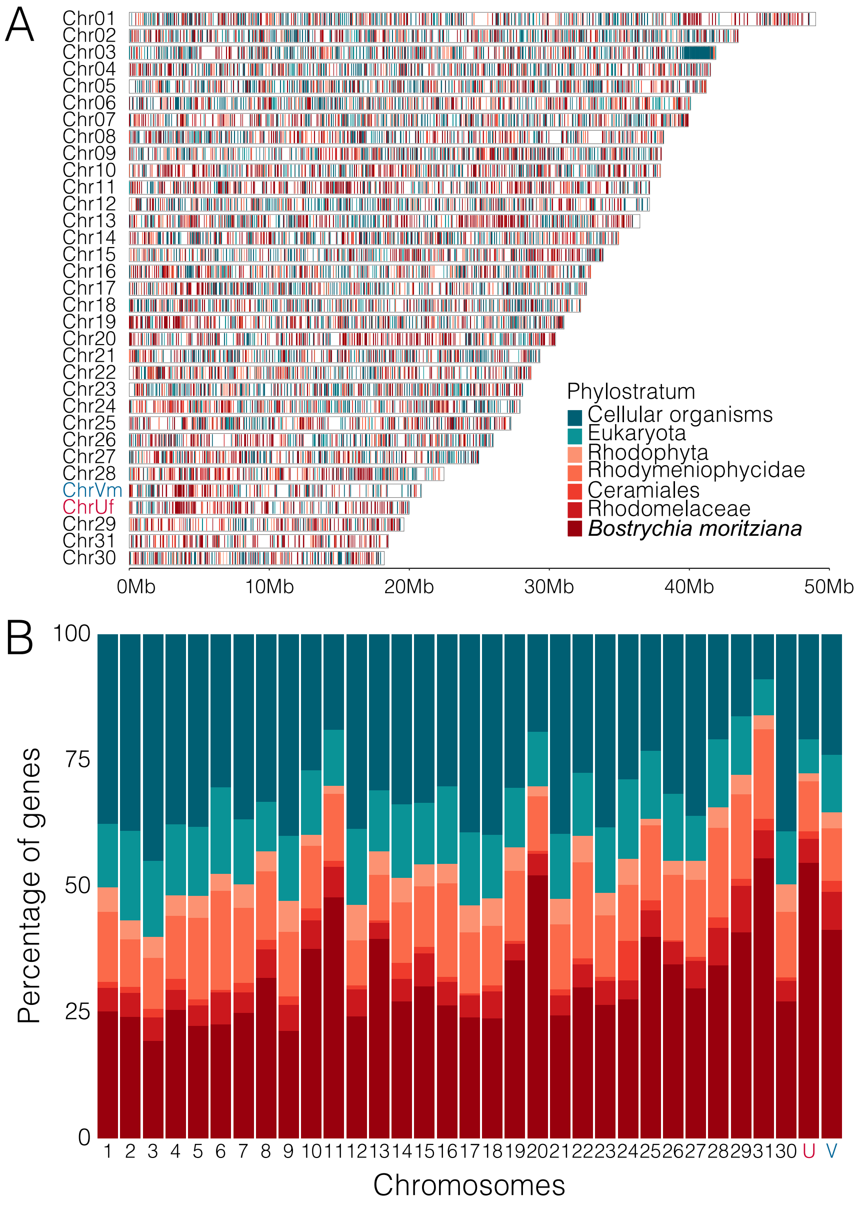


**Supplemental Figure 14. Gene age distribution across the *Bostrychia* genome.**
(**A**) Karyoplot displaying gene distribution across all chromosomes in *Bostrychia*. Genes are color-coded according to their assigned phylostrata as indicated in the legend. A scale bar indicating chromosome size (in Mb) is shown below the chromosomes. (**B**) Stacked bar charts represent the percentage of genes assigned to each phylostratum for each chromosome. The x-axis shows individual chromosomes, while the y-axis shows the proportion of genes belonging to different phylostrata. Colors correspond to gene age as indicated in the legend in panel A.

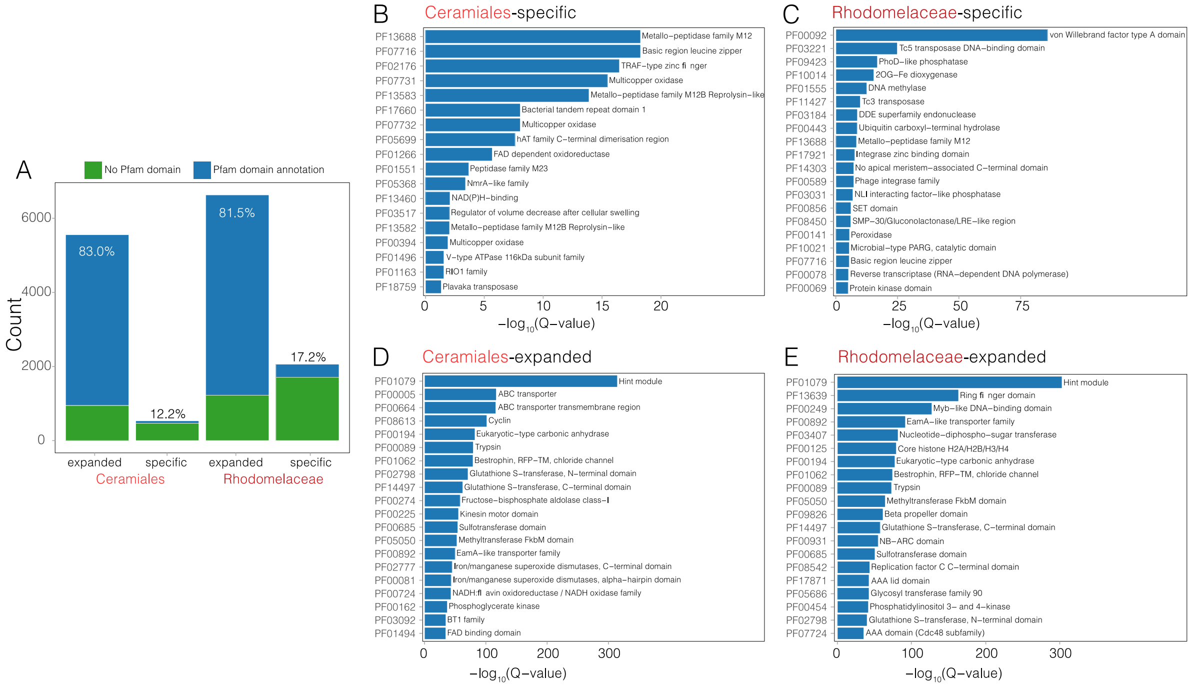


**Supplemental Figure 15. Annotated Pfam domains in *Bostrychia* genes.**
(**A**) Bar chart showing the number of genes in the expanded and specific gene families in the Ceramiales and Rhodomelaceae, with counts displayed on the y-axis. Green represents genes without an annotated Pfam domain, while blue indicates genes with an assigned Pfam domain. The percentage above each bar represents the proportion of genes with an annotated Pfam domain. (**B–E**) Results of the Pfam domain enrichment analysis for Ceramiales-specific (**B**), Rhodomelaceae-specific (**C**), Ceramiales-expanded (**D**), and Rhodomelaceae-expanded (**E**) gene families. The 20 most enriched Pfam domains are shown, with the x-axis displaying the -log$_{10}$(Q-value), where higher values indicate stronger statistical significance of Pfam domain enrichment. See **Supplemental Tab. 8** for the complete set of enriched Pfam domains.

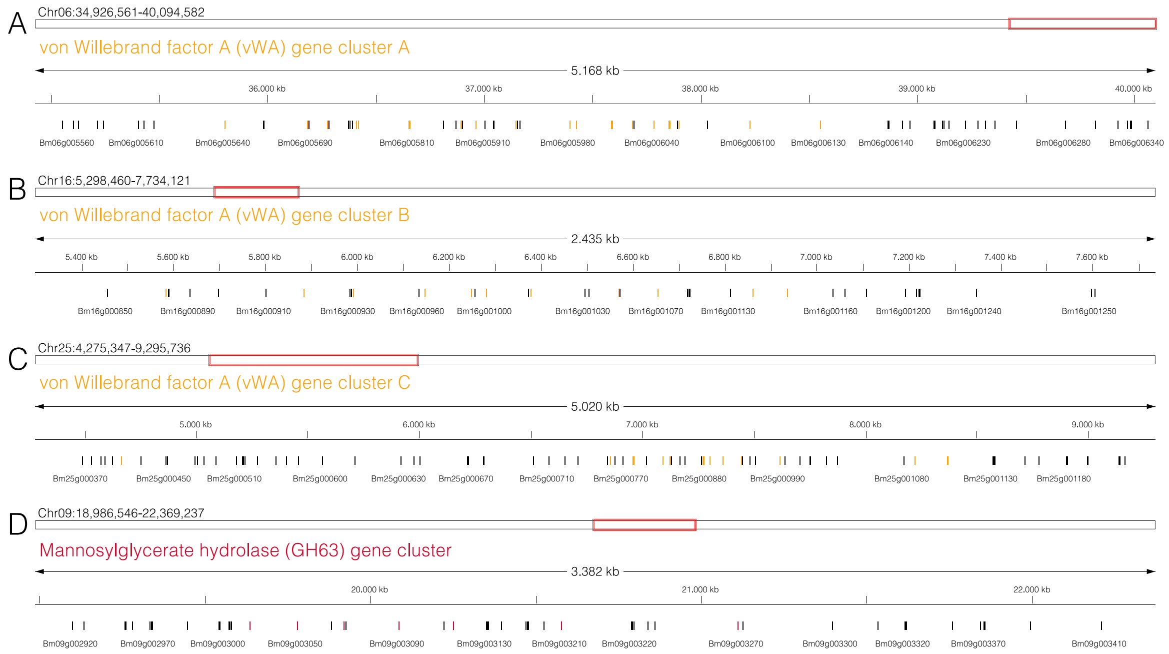


**Supplemental Figure 16. Genome browser view of selected gene clusters in *Bostrychia*.** Screenshots display four gene clusters across different chromosomes of *Bostrychia*. (**A-C**) Tandemly-repeated gene cluster of “von Willebrand factor A (vWA) domain” containing genes on chromosomes 6 (**A**), 16 (**B**) and 25 (**C**). Genes containing the vWA domain are shown in orange. (**D**) Gene cluster of seven mannosylglycerate hydrolases (GH63) on chromosome 9. Mannosylglycerate hydrolase genes are shown in red.

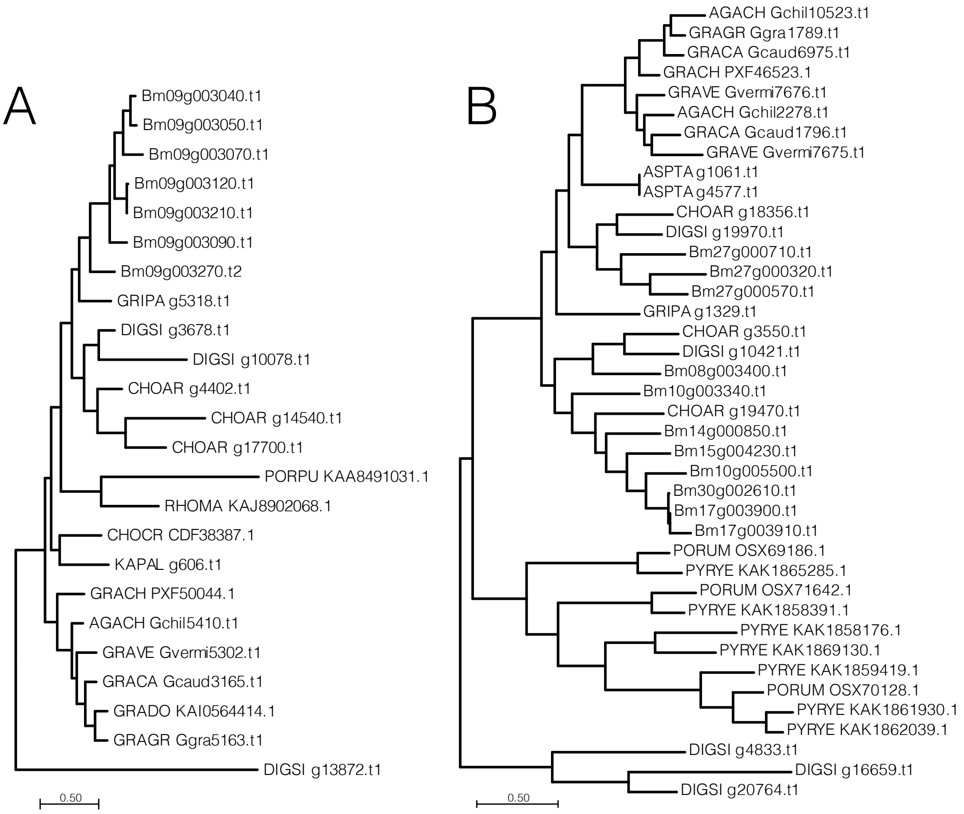


**Supplemental Figure 17. Unrooted phylogenetic trees of mannosylglycerate hydrolases and β-1,4-mannanases from red algae.**

Phylogenetic trees inferred for (**A**) mannosylglycerate hydrolase (GH63) and **(B**) β-1,4-mannanases (GH113) proteins from 20 red algal species. Predicted GTs and GHs were determined from annotated proteomes of 19 red algae species and the *Bostrychia* proteome.

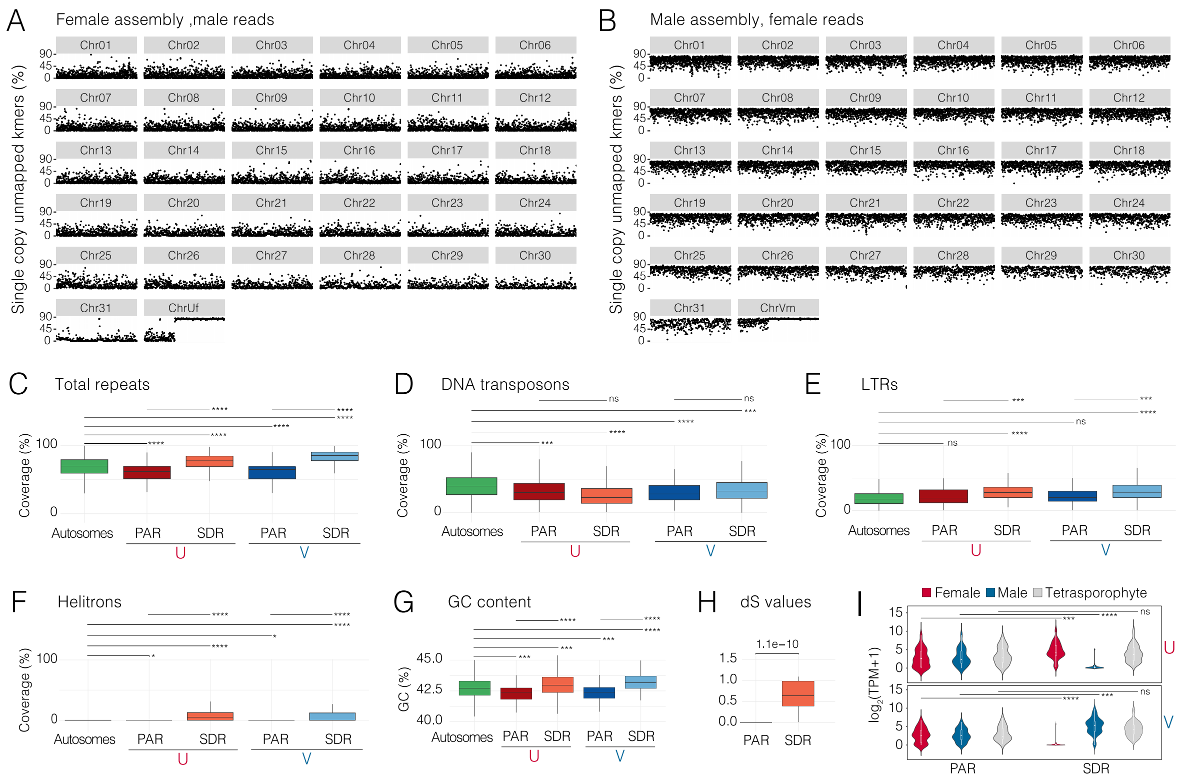


**Supplemental Figure 18. Identification and analysis of sex chromosomes in *Bostrychia*.**
(**A-B**) Dot plot showing the percentage of single-copy unmapped k-mers per chromosome in the female (**A**) and male gametophyte (**B**). The x-axes represent chromosome length while the y-axes indicate the proportion of unmapped k-mers. k-mers were mapped reciprocally to the fragmented assembly (50-kb chunks) of the opposing sex. Each dot represents the percentage of single-copy unmapped k-mers in a 50 kb-sized chunk. (**C-F**) Boxplots showing the percentage coverage of total repeats (**C**), DNA transposons (**D**), LTR elements (**E**), and Helitrons (**F**). (**G**) Boxplots displaying GC content percentages. The boxplots compare each category across autosomes (green), the female PAR (red), the female SDR (orange), the male PAR (dark blue) and the male SDR (light blue). (**H**) Boxplots comparing synonymous substitution rates (dS) of homologous PAR genes and SDR genes in the male and female gametophyte. (**I**) Violin plots showing the distribution of log2 transformed TPM+1 values of PAR and SDR genes on the U and V sex chromosomes in the female gametophyte (red), male gametophyte (blue) and tetrasporophyte (grey). Wilcoxon test results for statistical comparisons are shown above the boxplots and violin plots. * $p$-value < 0.05, ** $p$-value < 0.01, *** $p$-value < 0.001, ns = no significance.

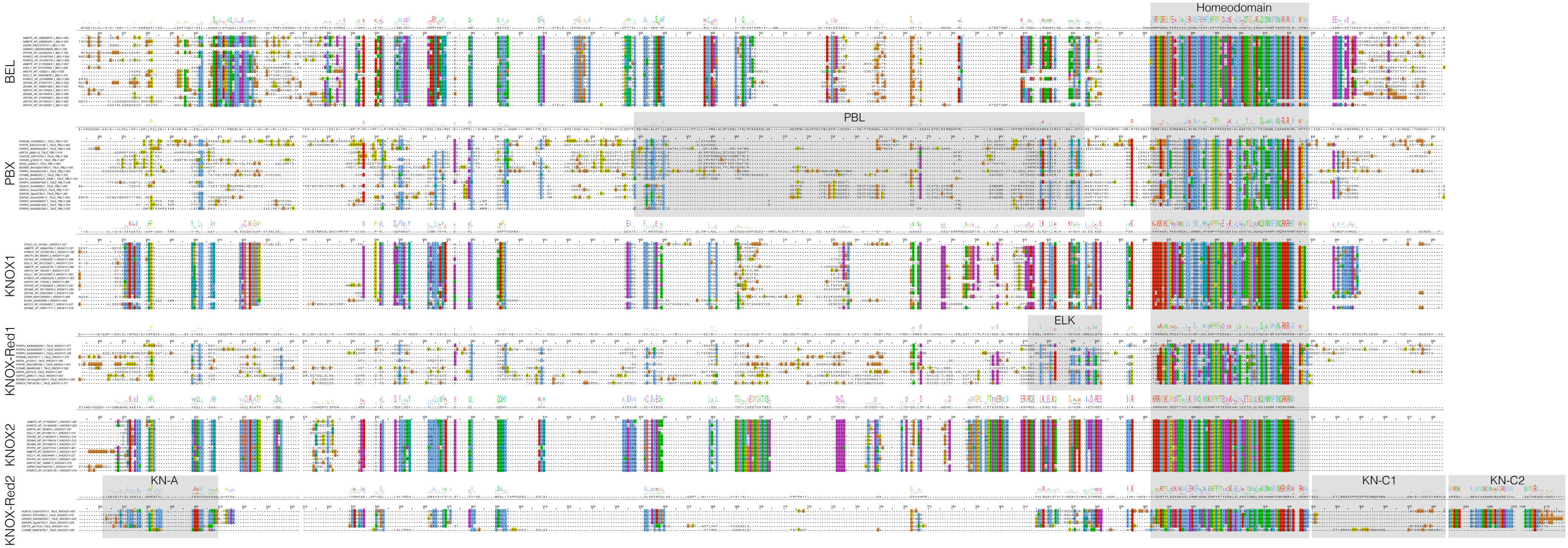


**Supplemental Figure 19. Alignment of TALE-HD transcription factors across the Archaeplastida.**
Alignment of TALE-HD protein sequences of selected red algae, plants, and streptophyte algae. The alignment is split into the red algal PBX, KNOX-Red1 and KNOX-Red2 classes using the plant BEL, KNOX1 and KNOX2 classes as comparison. The color-code refers to the conserved positions using the Clustal color scheme. The sequence logo represents conservation at each position. Grey boxes highlight conserved protein domains and sequence motifs as described previously[3].

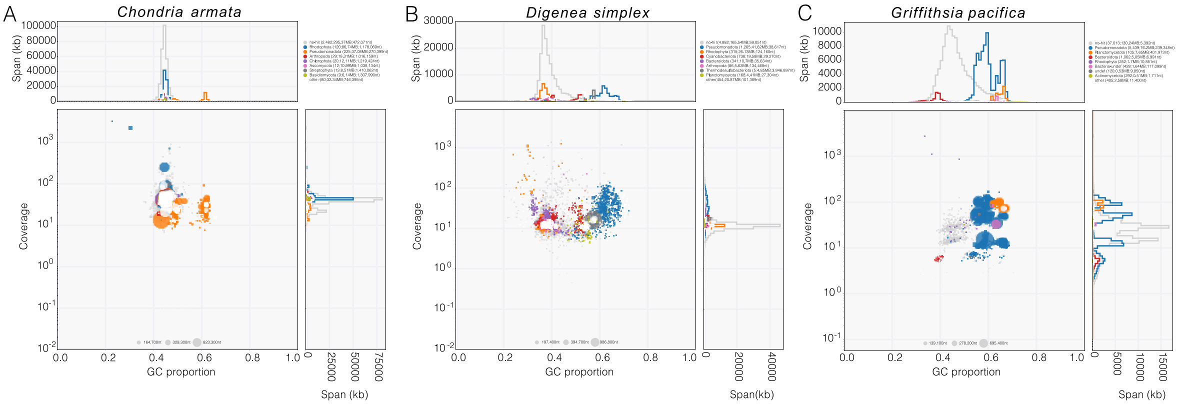


**Supplemental Figure 20. Blob plots of the *Chondria, Digenea* and *Griffithsia* draft assemblies.**
Blob plots displaying the taxonomic composition of contigs of the *Chondria* (**A**), *Digenea* (**B**) and *Griffithsia* (**C**) genome assemblies. Each point represents an individual contig, with its position on the x-axis determined by GC content and sequencing coverage on the y-axis. Contigs are color-coded according to their assigned taxonomy. The size of each point corresponds to the contig length. The span in kb of each taxonomic group for each GC proportion is shown above and next to each dot plot.
